# An atlas of *Caenorhabditis elegans* chemoreceptor expression

**DOI:** 10.1101/222570

**Authors:** Berta Vidal, Ulkar Aghayeva, Haosheng Sun, Chen Wang, Lori Glenwinkel, Emily Bayer, Oliver Hobert

## Abstract

One goal of modern day neuroscience is the establishment of molecular maps that assign unique features to individual neuron types. Such maps provide important starting points for neuron classification, for functional analysis and for developmental studies aimed at defining the molecular mechanisms of neuron identity acquisition and neuron identity diversification. In this resource paper, we describe a nervous system-wide map of the potential expression sites of 244 members of the largest gene family in the *C. elegans* genome, rhodopsin-like (class A) GPCR chemoreceptors, using classic *gfp* reporter gene technology. We cover representatives of all sequence families of chemoreceptors GPCRs, some of which were previously entirely uncharacterized. Most reporters are expressed in a very restricted number of cells, often just in single cells. We assign GPCR reporter expression to all but two of the 37 sensory neuron classes of the sex-shared, core nervous system. Some sensory neurons express a very small number of receptors, while others, particularly nociceptive neurons, co-express several dozen GPCR reporter genes. GPCR reporters are also expressed in a wide range of inter- and motorneurons, as well as nonneuronal cells, suggesting that GPCRs may constitute receptors not just for environmental signals, but also for internal cues. We observe only one notable, frequent association of coexpression patterns, namely in one nociceptive amphid (ASH) and two nociceptive phasmid sensory neurons (PHA, PHB). We identified GPCRs with sexually dimorphic expression and several GPCR reporters that are expressed in a left/right asymmetric manner. We identified a substantial degree of GPCR expression plasticity; particularly in the context of the environmentally-induced dauer diapause stage when one third of all tested GPCRs alter the cellular specificity of their expression within and outside the nervous system. Intriguingly, in a number of cases, the dauer-specific alterations of GPCR reporter expression in specific neuron classes are maintained during postdauer life and in some case new patterns are induced post-dauer, demonstrating that GPCR gene expression may serve as traits of life history. Taken together, our resource provides an entry point for functional studies and also offers a host of molecular markers for studying molecular patterning and plasticity of the nervous system.

**AUTHOR SUMMARY:** Maps of gene expression patterns in the nervous system provide an important resource for neuron classification, for functional analysis and for developmental studies that ask how different neurons acquire their unique identities. By analyzing transgenic gfp reporter strains, we describe here the expression pattern of 244 putative chemosensory receptor-encoding genes, which constitute the largest gene family in *C.elegans*. We show that, as expected, chemoreceptor expression is enriched in chemosensory neurons but it is also expressed in a wide range of interneurons, motorneurons, as well as non-neuronal cells, suggesting that putative chemosensory receptors may not just sense environmental signals but also internal cues. We find that each chemoreceptor is expressed in a few neuron types, often just one, but each neuron type can express a large number of chemoreceptors. Interestingly, we uncovered that chemoreceptor expression is remarkably plastic, particularly in the context of the environmentally-induced dauer diapause stage. Taken together, this molecular atlas of chemosensory receptors provides an entry point for functional studies and offers a host of markers for studying neuronal patterning and plasticity.

## INTRODUCTION

Molecular markers selectively expressed in individual neuron types represent invaluable tools to understand how cellular diversity in a nervous system is genetically encoded. Molecular markers that are constitutively and invariably expressed throughout the life of a specific neuron type provide static views of neuronal identity and hence provide entry points to study how invariable identity features are acquired during neuronal differentiation [1]. In contrast, some molecular features of a neuron display a remarkable plasticity in that their expression may be regulated by neuronal activity or in response to specific environmental cues. Such genes serve as markers to understand the nature of the gene regulatory programs that govern such dynamic features of a neuron. We reasoned that a significant expansion of the expression analysis of chemosensory G-protein-coupled receptors (GPCRs), initiated more than 20 years ago [2] using *gfp*-based reporter gene technology [3], may yield a significantly expanded resource of molecular markers that may label various aspects of neuronal identity and neuronal plasticity in the *C. elegans* nervous system.

Animal genomes encode five major classes of GPCRs, of which the rhodopsin class (or “class A”) is the largest class [4, 5](Table 1). Rhodopsin class GPCRs can be subdivided into phylogenetically deeply conserved neurotransmitter receptors (neuropeptides, acetylcholine, biogenic amines) as well as non-conserved, chemosensory-type GPCRs (from here on referred to as “csGPCRs”)(Table 1). The csGPCRs have independently expanded in distinct animal phyla where they serve to respond to diverse, physiologically relevant external and, supposedly, internal cues [4, 6, 7]. The genome of the nematode *C. elegans* encodes an exceptionally large battery of chemosensory-type csGPCRs composed of 1,341 protein-coding genes (Table 2)[2, 7, 8], a remarkable number given the small size of its nervous system (302 neurons constituting 118 anatomically defined neuron types)[9]. These csGPCRs have been subdivided by sequence into families and superfamilies, as summarized in Table 2 [2, 7].

**Table 1:**
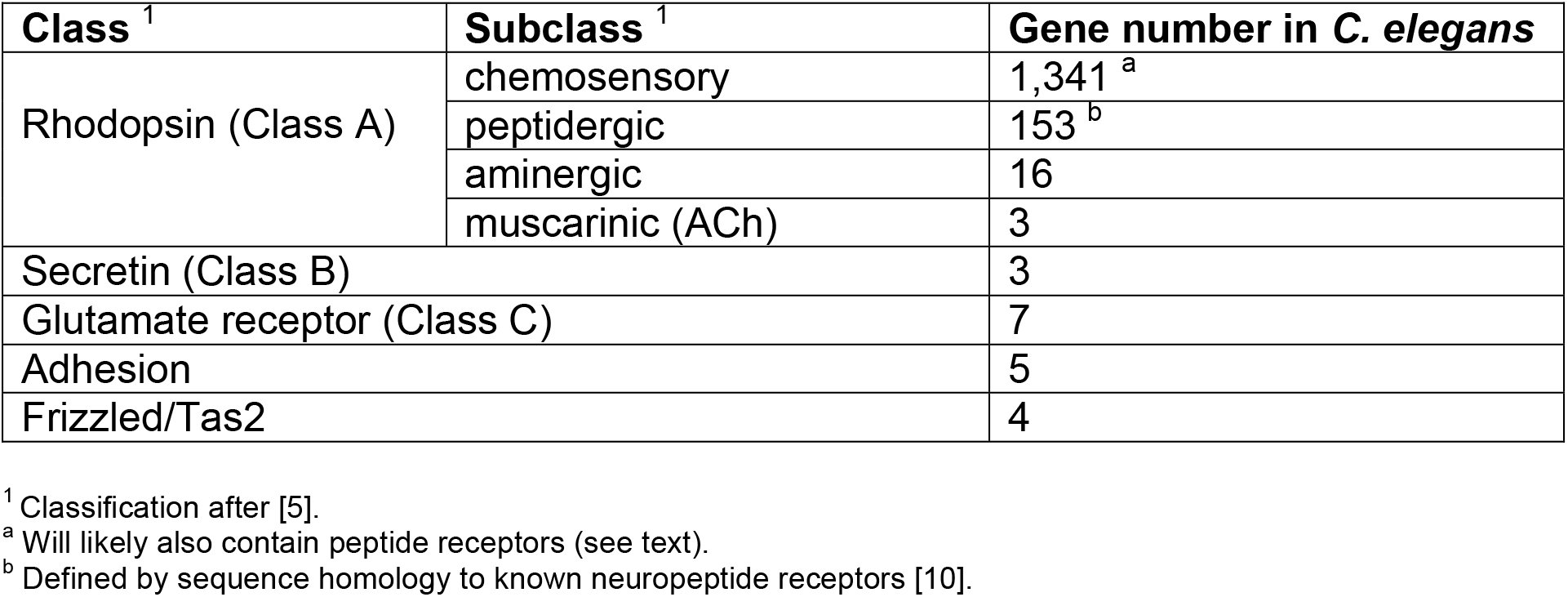
The five classes of GPCRs in animal genomes and their representation in *C.elegans*. Modified from [10].

| **Class**<sup>1</sup> | **Subclass**<sup>1</sup> | **Gene number in *C. elegans*** |
| --- | --- | --- |
| Rhodopsin (Class A) | chemosensory | 1,341<sup>a</sup> |
|  | peptidergic | 153<sup>b</sup> |
|  | aminergic | 16 |
|  | muscarinic (ACh) | 3 |
| Secretin (Class B) |  | 3 |
| Glutamate receptor (Class C) |  | 7 |
| Adhesion |  | 5 |
| Frizzled/Tas2 |  | 4 |

<sup>1</sup> Classification after [5].
<sup>a</sup> Will likely also contain peptide receptors (see text).
<sup>b</sup> Defined by sequence homology to known neuropeptide receptors [10].

**Table 2:**
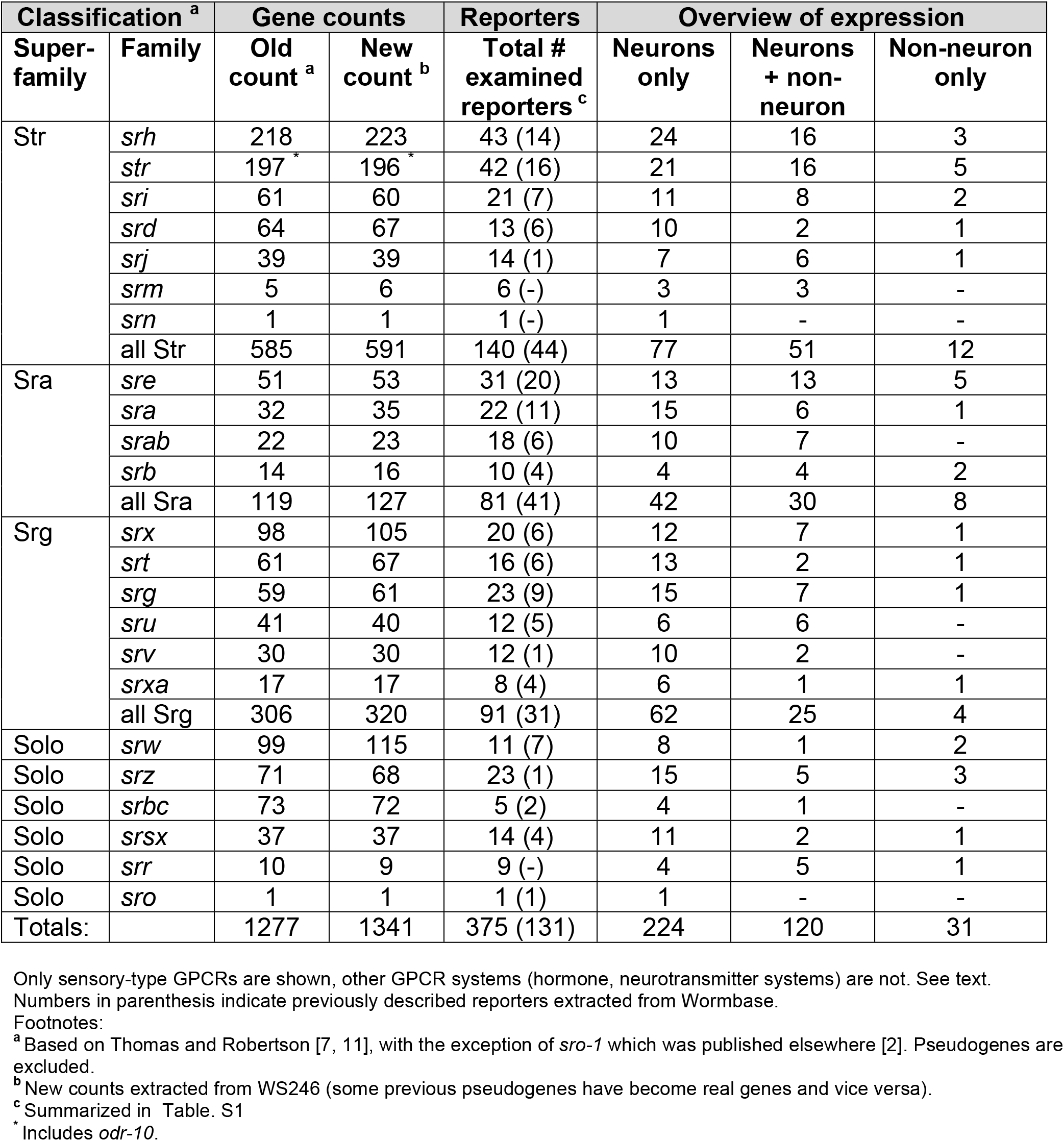
Overview of GPCR reporters and expression

Wormbase contains expression data for 131 csGPCRs, however for only 76 of them the expression site has been defined with single cell resolution (S1 Table). The majority of these 76 reporters revealed expression in chemosensory neurons [2]. Functional studies have linked a small subset of these receptors to the sensation of specific environmental or pheromonal cues [12–21], but in the absence of concerted de-orphanization efforts like those seen in other organisms [22, 23], the number of receptors with assigned ligands is still remarkably low.

Intriguingly, a subset of the previously characterized csGPCR genes were also expressed in non-sensory neurons [2, 24–28] suggesting that they may also function as receptors of internal ligands of unknown identity. Providing some hints to the identity of these ligands, one csGPCR subclass, encoded by the *srw* genes, displays sequence similarities to peptide receptors [11, 29]. The expression of csGPCRs in interneurons also prompted efforts to identify the function of some of these genes. Even though its ligand remains unknown, AIY-expressed *sra-11* was found to be involved in the associative learning paradigm, olfactory imprinting [30], while *sra-13* acts in the vulva to control vulval development, which is affected by food signals [26].

In spite of the relative paucity of known ligands, the previously published expression patterns of csGPCRs provided molecular indicators for a number of intriguing and generally very poorly understood nervous system features: (1) the expression pattern of the GPCR gene *str-2* revealed a left/right asymmetry in the two AWC olfactory neurons [31]; this lateralization phenomenon was later found to be required for olfactory discrimination [32] and spurred a host of studies aimed at revealing how this left/right asymmetry is developmentally programmed [33]. (2) The expression of several csGPCRs revealed a remarkable plasticity in response to changes in the environment. For example, expression of *srd-1* and *str-2* and *str-3* changes in ASI neurons in response to dauer pheromone [34], and expression of *srh-34* and *srh-234* in ADL is different in fed *versus* starved animals [35]. Using these dynamic reporter gene patterns, mechanisms controlling csGPCR plasticity have been elucidated [35, 36]. (3) The csGPCR genes *srd-1, srj-54* and *odr-10* have been found to be expressed in a sexually dimorphic manner in sex-shared sensory neurons, suggesting that sexual identity impinges on sensory perception [2, 37, 38].

In this resource paper, we examined the expression of 244 reporter transgenes that monitor expression of previously uncharacterized csGPCR genes (for simplicity, from here on referred to as “GPCRs”. Our explicit goal in this analysis was to (1) generate more neuronal identity markers, (2) test the hypothesis that many more sensory neurons may be lateralized, (3) identify more markers of neuronal plasticity, (4) identify more markers of sexual dimorphism and (5) examine the extent of expression in non-sensory and non-neuronal cells (suggesting roles as receivers of internal signals). Based on the molecular classification of csGPCRs into defined families, we were also interested in determining whether the expression of specific subfamilies – particularly those whose expression has not previously examined – may reveal specific common themes (*i.e*., patterns of co-expression or expression in specific cells) that may provide a hint to their function. We synthesize our findings with those of previous expression pattern analyses to carve out a number of general features of csGPCR expression patterns.

## MATERIALS AND METHODS

### Mutant Strains

Strains were maintained by standard methods [39]. Mutant alleles used in this study were: *pha-1(e2123)* [40], *him-5(e1490)* [41], *unc-43(n1186lf)* [42], *unc-43(n498gf)[43]* and *nsy-5(ky634)* [44].

### Reporter and transgenic strain generation

GFP reporters were generated using a PCR fusion approach [45] and injected without being subcloned. Genomic fragments were fused to the GFP coding sequence, which was followed by the *unc-54 3*’ untranslated region. A list of primers for all constructs can be found in the Supplementary Methods. Amplicons were injected at 50ng/ml with the *pha-1* rescuing plasmid (pBX) as a co-injection marker (50ng/ml). Reporters were injected into a *pha-1(e2123)* or *pha-1(e2123);him-5(e1490)* mutant background strain [40], resulting in transgenic arrays with little mosaicism. For each construct 2 independent lines were scored. Reporter strains provided by the Vancouver Consortium were generated as described [46]. Further details and primer sequences used by the Vancouver Consortium can be found at http://www.gfpworm.org. A list of all reporter strains generated by us or provided by the Vancouver Consortium can be found in the Supplementary Methods.

### Microscopy

Worms were anesthetized using 100mM sodium azide (NaN_3_) and mounted on 5% agarose on glass slides. Images were acquired using an automated fluorescence microscope (Zeiss, AXIO Imager Z.2). Acquisition of several z-stack images (each ~1 mm thick) was performed with the ZEN 2 pro software. Representative images are shown following max-projection of Z-stacks using the maximum intensity projection type. Image reconstruction was performed using Fiji software [47].

### Neuron identification

Neurons were identified either by labeling subsets of sensory neurons with DiD (Thermo Fisher Scientific) or by crossing reporter transgenes with landmark reporter strains in which known neuron types are labeled with a red fluorescent reporter. For dye filling, worms were washed with M9, incubated with DiD (1:500) in M9 for 1 hour at room temperature, washed 3 times with M9, and plated on agar plates coated with food for 1-3 hours before imaging. Red fluorescent reporter strains used for cell identification are: *otIs263[ceh-36p::TagRFP, rol-6(su1006)], vyIs51[str-2p::2xnls::TagRFP; ofm-1p::DsRed]*[48], *otIs518[eat-4^Fosmid^::sl2::mCherry::h2b]*[49], *otIs544[cho-1^Fosmid^::sl2::mCheriy::h2b]*[50], *otIs564[unc-47^Fosmid^::sl2::mCherry::h2b]*[51], *otIs612[flp-18p::NLG-1::GFP11, gpa-6p::NLG-1:::GFP1-10, flp-18p::mCherry, nlp-1p::mCherry], hdIs30[glr-1p::DsRed], otIs521[eat-4prom8::tagRFP; ttx-3::gfp]*.

### Hierarchical clustering of neurons by GPCR reporter expression

Clustering was performed on binary expression data from 272 neuron-expressed GPCR reporters for which we had cell ID information. Expression data was from our own analysis and available data from <u>wormbase.org</u> [52]. Only positive neuronal cell ID information per GPCR reporter was included in the binary expression matrix with no distinction between the absence of expression and unknown expression per neuron. Data were clustered using the R pvclust package (<u>https://cran.r-project.org/web/packages/pvclust/pvclust.pdf</u>) [53] using the euclidean distance metric with average linkage, bootstrap 1000, and relative sample size ranging from a proportion of 0.5 to 1.4 of the original sample size. The relative proportion was incremented by 0.1 for each bootstrap resampling. Bootstrap Probability value (BP) and Approximately Unbiased p-values (AU) are derived from the multiscale-multistep bootstrap resampling. AU support values > 95 indicate well-supported clusters and should be considered when evaluating dendrogram cluster relationships. Alternative distance and linkage methods showed clustering of the PHA, PHB and ASH neurons in all cases (42 out of 84 cases had strong support with AU/BP >95).

### Upstream intergenic distances and intron length calculations

GPCR upstream intergenic regions and intron lengths were extracted from *C. elegans* exon coordinates, version WS220 using a python script. Non-coding RNA exons were excluded from the intergenic distance calculations so that intergenic distances represent the nucleotide sequence distance between coding genes. The average intron length per gene was calculated by summing the intron sequence lengths for each gene and dividing by the total number of introns. Average intron lengths for genes with multiple isoforms were calculated for each isoform and then averaged, resulting in one average intron length per gene.

### Generation of dauers and analysis of changes in expression

To analyze GPCR reporter gene expression in dauers, mixed populations of respective strains were allowed to exhaust food for 5-7 days at 20°C. Dauers were isolated from starved plates by treatment with 1% SDS for 30 min and imaged within 1-2 hours of isolation. The cellular identity of expression changes in dauers were confirmed with red landmark strains, as mentioned above.

## RESULTS

### Selection of csGPCRs for expression analysis and method of analysis

We chose to examine csGPCR expression patterns using gfp-based reporter gene technology, the standard tool of gene expression analysis in *C. elegans* [3, 54]. The obvious shortcoming of this technology is that reporter genes may not capture the full *cis*-regulatory content of the respective GPCR-encoding locus, but as we will describe in more detail below, most GPCR-encoding loci are compact with small intergenic regions and introns. We emphasize that our approach is not necessarily aimed at identifying the complete set of cells expressing a GPCR, but, following ample precedent, is rather aimed at identifying novel and informative patterns of expression, as incomplete as these patterns may be.

We utilized two sources of csGPCR reporters. A consortium at the University of British Columbia (Vancouver) has generated a valuable, large panel of reporters for 1886 genes in the *C. elegans* genome [46]. However, the site of expression of these reporters has not been determined with single cell resolution in the nervous system. We obtained 100 reporters from this collection that targeted GPCR loci and for every reporter that produced a stable pattern of expression, we undertook a detailed analysis of their sites of expression in the nervous system.

In addition to these 100 reporter genes, we generated 144 of our own reporter genes. We adhered to the following principles in the choice of genes and design of reporters: First, we aimed to cover all 23 classes of chemoreceptor genes defined by Thomas and Robertson [7](Table 2). Using phylogenetic trees assembled by Thomas and Robertson, we sampled each gene family evenly, generally avoiding the examination of close sequence paralogues, which we anticipated to reveal similar expression patterns.

Our own reporters mostly contain all 5’ intergenic regions fused to *gfp* and contain at most 4 kb of sequence. The rationale behind this choice lies in the overall organization of GPCR loci (summarized in S1 Fig). 89% of the ~1,300 csGPCR loci contain 5’ intergenic regions of less than 4kb. We chose all of our samples from this pool and the reporters generated by us capture the full intergenic region. The reporters from the Vancouver consortium contain about 3 kb of 5’ intergenic region at most [46]. Furthermore, csGPCR loci tend to have small introns (average size 432 bp; almost half of them <200 bp; S1 Fig), indicating that relatively little *cis*-regulatory information resides in these introns, which provided the basis for our focus on intergenic regions. For some genes with very short upstream intergenic regions (less than 500 bp) we included the first intron (if this was 300 bp or larger) in order to increase the regulatory space contained in the reporters. The coordinates for all reporter constructs can be found in the Supplementary Material.

Sites of expression within the nervous system were determined mainly for those reporters with most robust expression and was based on stereotyped cell position, cellular and process morphology and co-labeling with either DiD (which labels a subset of sensory neurons) or by crossing with landmark strains in which specific neuron types are labeled with a red fluorescent protein (see Material and Methods). All cell identification was initially done in young adult hermaphrodite animals. As we will describe in detail later, a number of these reporter strains were also subjected to analysis at different stages, under different conditions and in the two different sexes.

### GPCRs are expressed in restricted patterns within and outside the nervous system

In our ensuing description of expression patterns of reporter genes, we summarize the expression observed with the previously described reporters, as well as the additional reporters analyzed by us. All of our expression analysis is summarized in a tabular form in S1 Table. Three overall features of the 375 csGPCR reporters are immediately apparent (Fig 1): first, 92% of analyzed reporters are expressed in the nervous system; second, expression is not restricted to the nervous system: 33% of the reporters are expressed both within and outside the nervous system and 8% are expressed exclusively in non-neuronal cells, and third, the vast majority of csGPCR reporters are expressed in very restricted number of cells (Fig 1A,B). Of the neuronally expressed reporters, 24% are expressed in single neuron pairs, 27% in 2 neuron pairs, 26% in 3-4 neuron pairs, 19% in 5-10 neuron pairs and the remaining 4% in more than 10 neuron pairs.

**Fig 1.**
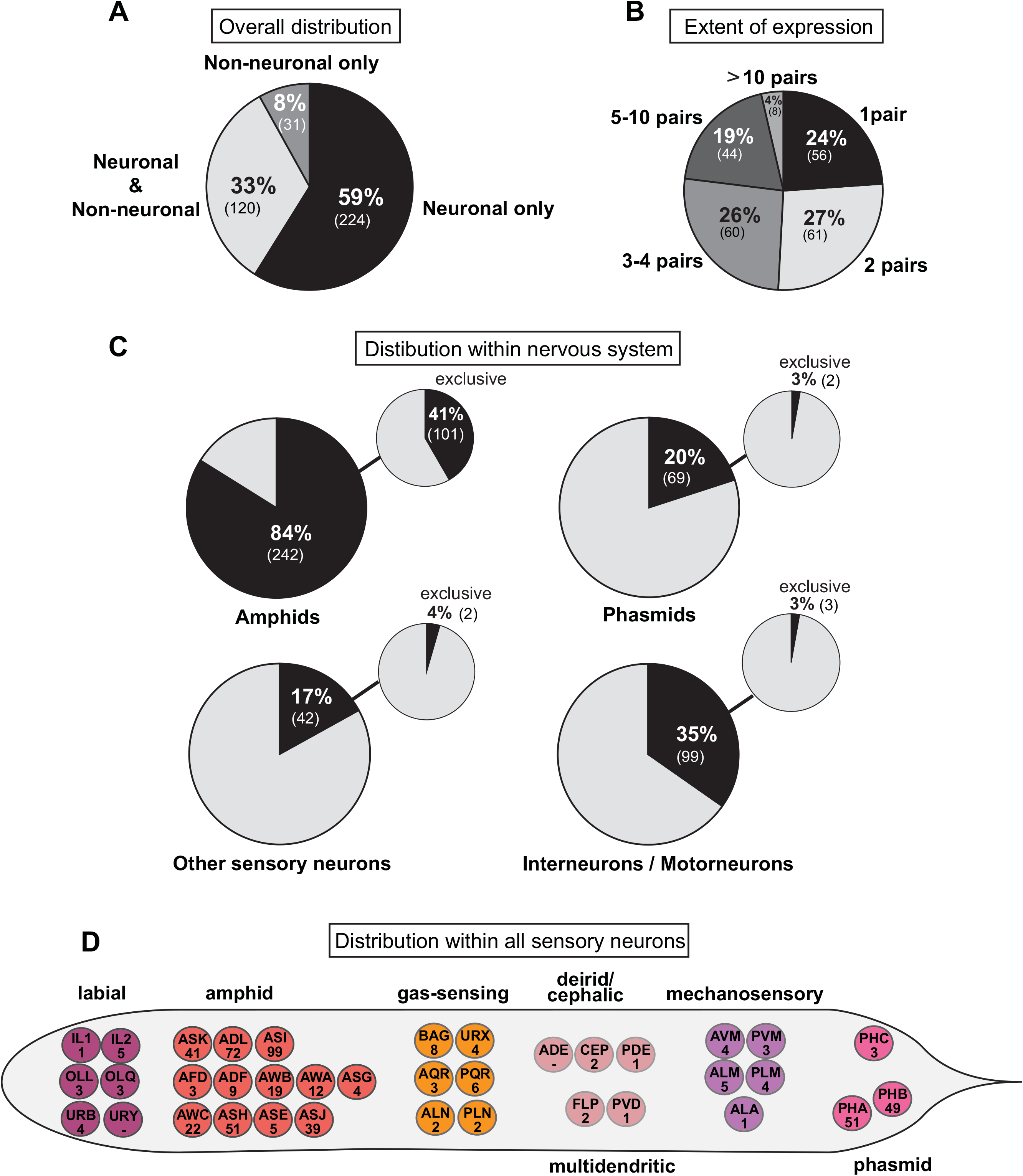
Summary of GPCR reporter expression patterns. (**A**) Overall tissue distribution of reporter expression patterns. Pie chart showing percentage of GPCR reporters expressed exclusively in neurons, in neurons and other cells types and exclusively in non-neuronal tissues. Numbers in parenthesis represent the absolute number of reporters in each category. (**B**) Extent of reporter expression within the nervous system. Pie chart showing percentage of neuronal reporters expressed in 1 neuron pair, 2 pairs, 3-4 pairs, 5-10 pairs or more than 10 pairs. Numbers in parenthesis represent the absolute number of reporters in each category. (**C**) Distribution of reporter gene expression within the nervous system. Pie charts showing percentage of GPCR reporters expressed in amphid neurons, phasmid neurons, other sensory neurons and inter- or motorneurons. Small pie charts on the upper right represent the percentage of reporters exclusively expressed in amphids, phasmids, other sensory neurons and inter- or motorneurons. Numbers in parenthesis show the absolute number of reporters in each category. (**D**) Distribution within all sensory neurons. Worm schematics showing the absolute number of GPCR reporters found to be expressed in each sensory neuron type. PHC is a phasmid neuron by name only (“PH”). See Table S2 for a list of GPCR gene names expressed in the sensory neurons shown here.

Expression outside the nervous system will be described in a later section. Within the nervous system, expression is most prominent in sensory neurons (Fig 1C). 84% of the reporters are expressed in amphid sensory neurons (which are made up of 12 pairs of neurons), 20% in phasmid sensory neurons (made up of 2 pairs of neurons, PHA and PHB), and 17% in other sensory neurons. We find that every sensory neuron, except for URY and ADE neurons, expresses at least one GPCR (Fig 1D; Table S2). The number of GPCRs expressed in a given neuron shows a striking range. The ASI neuron expresses an impressive 99 GPCR reporters. After ASI, the nociceptive neurons ADL and ASH together with the phasmid neurons PHA and PHB are the sensory neurons with higher number of GPCRs, expressing 72, 51, 51 and 49 reporters respectively. Outside the amphid and phasmid neurons, the number of reporters expressed in sensory neurons dramatically drops, with all other sensory neurons expressing less than 10 GPCRs, in some cases only a single GPCR (Fig 1; Table S2). Of course, it needs to be kept in mind that we only consider expression of a fraction of the csGPCR loci and hence each of these total numbers is expected to increase by several fold once all csGPCR expression patterns are identified.

24% of the GPCR reporters for which we have information about neuron numbers are exclusively expressed in a single neuron class and in all these cases, the neuron class is a sensory neuron class (Fig 2; Table S3). In total, however, only nine sensory neurons express single-neuron specific GPCRs. The most striking one of them is the ADL nociceptive neuron, which expresses 23 single neuron-specific GPCR reporters (and an additional 49 GPCR reporters expressed in additional neurons). The ADL-expressed, single neuron-specific GPCRs do not fall into a specific GPCR subfamily but rather cover 7 distinct families. A small subset of the single neuron type specific GPCRs are expressed outside the nervous system as well (genes with asterisk in Fig 2A). This may indicate that these receptors do not detect external cues, but rather sense internal signals.

**Fig 2.**
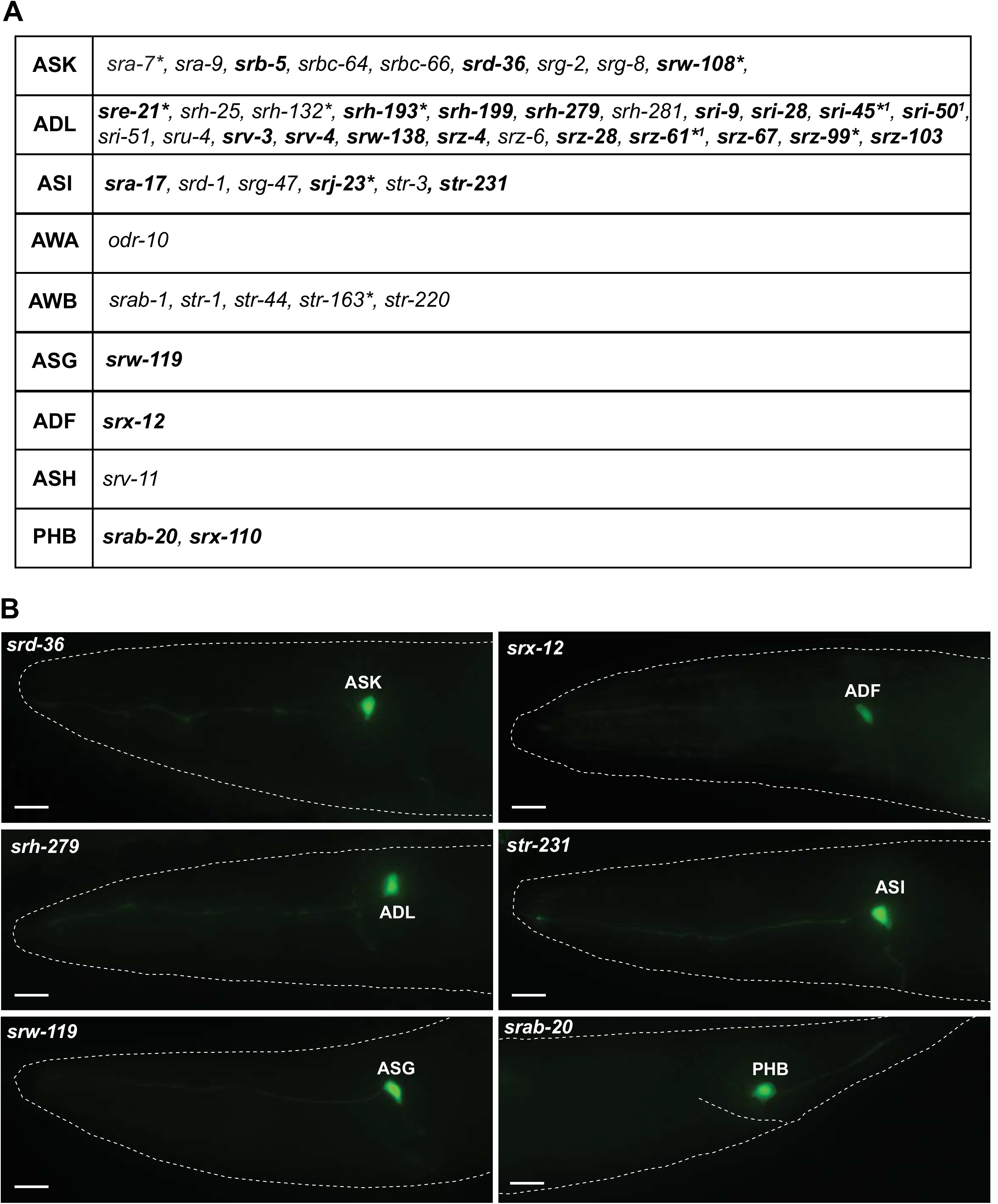
GPCR reporters expressed in single sensory neuron classes. (**A**) Table showing all GPCR reporters expressed in a single neuron class. Genes in bold are newly identified in this paper. Genes in non-bold were previously described (data extracted from <u>www.wormbase.org</u>). * Reporter is also expressed in some non-neuronal tissue (for details see S1 Table).^1^ N. Masoudi, S. Finkelstein and O. Hobert, in preparation. (**B**) Representative examples of reporters expressed in a single neuron class identified in this study. Scale bars, 10μm.

Notably, expression of the csGPCR reporter collection is clearly not restricted to sensory neurons. A striking 35% of the csGPCR reporters are expressed in inter- and motorneurons (Fig 1, Fig 3; Table 3; S1 Table). There is no unifying feature of the inter- or motorneurons that express GPCR reporters. They range from ventral cord motor neurons to head interneurons, and to command interneurons in the ventral cord. One interneuron, PVT, displays a very large number of expressed csGPCR reporters (57 different reporters); however, PVT expression is generally observed in an unusually large amount of reporter genes and may, like posterior gut expression, be a reporter gene artifact that relies on cryptic regulatory elements in the reporter.

**Fig 3.**
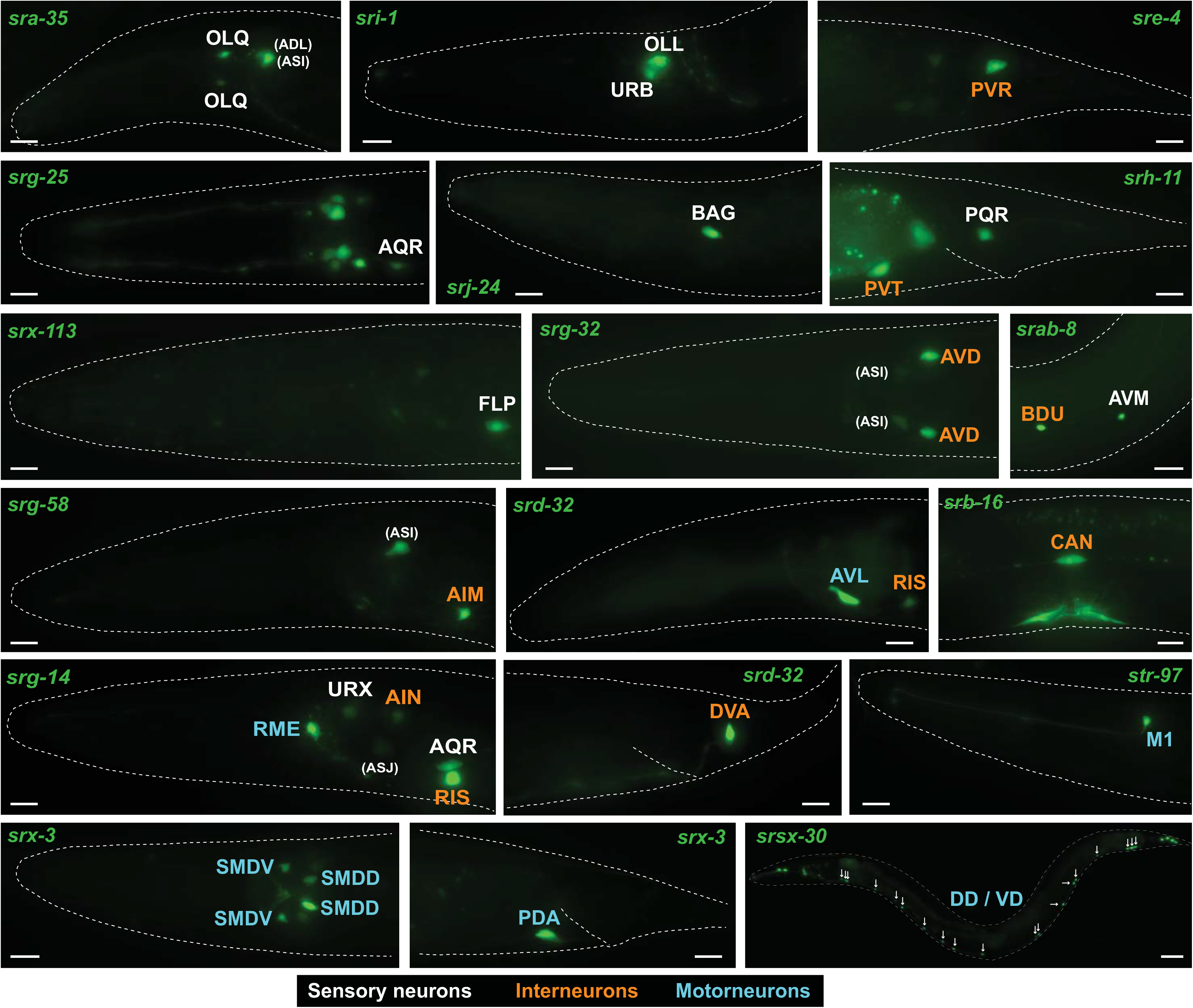
GPCR reporters expressed in non-amphid/non-phasmid sensory neurons, interneurons and motorneurons. Examples of GPCR reporters expressed in sensory neurons that are not amphids or phasmids (white font), interneurons (orange font) and motorneurons (blue font). Most examples represented here are from neurons classes that were not previously shown to express any sensory GPCR. Amphid neurons are shown in parenthesis. All scale bars,10μm, except *srsx-30* which is 30μm. See Table 3 for a complete summary of GPCR reporters expressed in inter- and motorneurons.

**Table 3:**
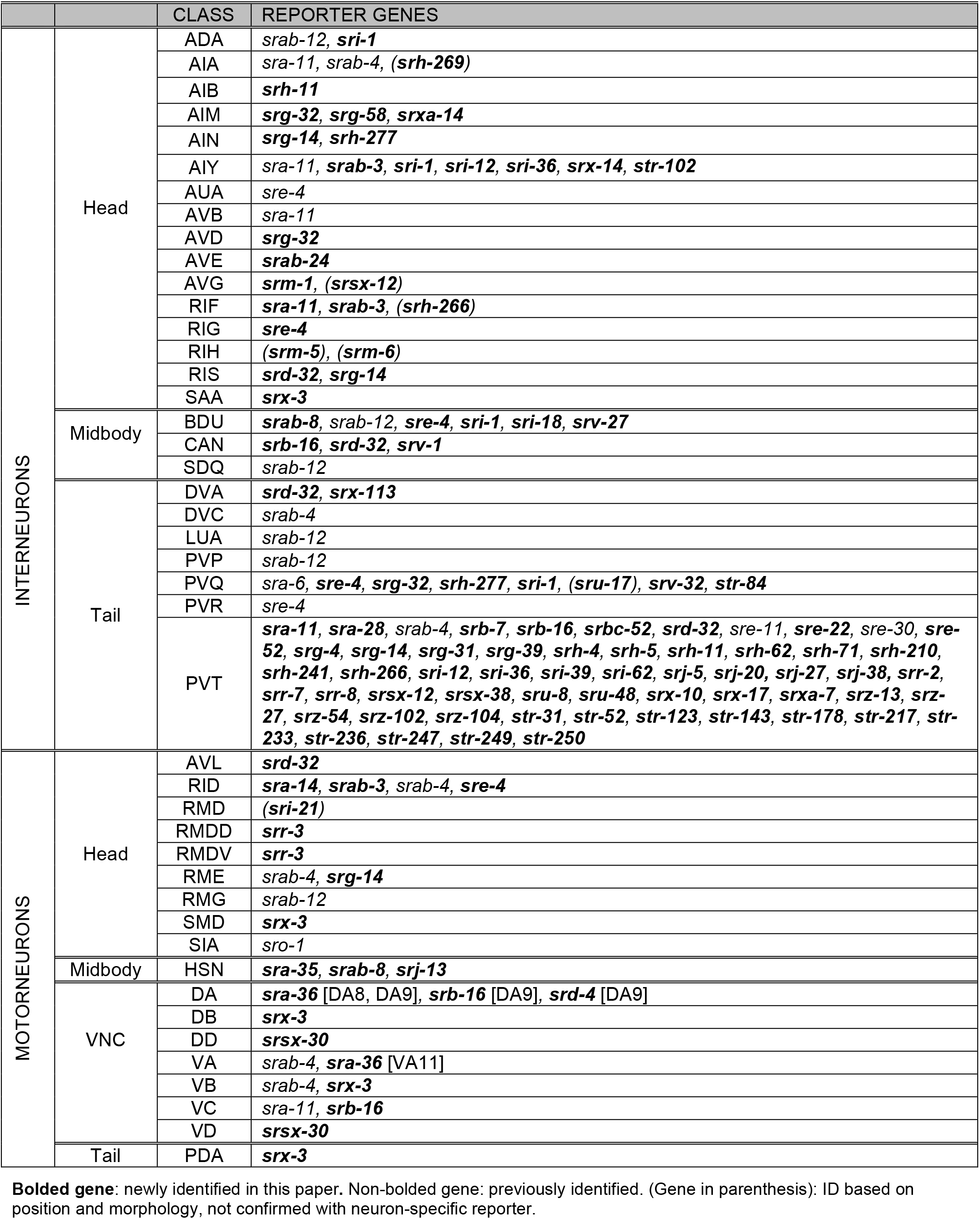
Non-sensory neurons expressing GPCR reporter

97% of inter and/or motorneuron-expressed csGPCR reporters are also expressed in sensory neurons so only 3% of them show expression exclusively in inter or motorneurons. In light of the inter/motorneuron expression of csGPCR reporters, one can imagine that csGPCR reporters that are expressed in sensory neurons may actually not function as receptors for external sensory cues, but may rather function as they likely do in inter/motorneurons, *i.e*. as receptors of internal signals.

We asked whether csGPCR expression profiles cluster by neuron type. To this end, we undertook unsupervised hierarchical clustering of expression profiles. The bootstrap value for most associations was very weak with two exceptions: csGPCR reporters are often coexpressed in the two tail phasmid neuron classes PHA and PHB (AU/BP>95) and expression in either or both of the phasmid neurons is associated with the expression in the head neuron ASH (AU/BP>95) (Fig 4). These associations are striking since all these 3 neuron classes are nociceptive neurons that respond to some similar cues and integrate sensory inputs from the head and tail [55, 56] and that directly innervate command interneurons involved in reversal behavior [9]. While GPCRs expressed in these neurons are likely involved in sensing nociceptive cues, it is notable that these co-expressed csGPCR came from a widely distinct sets of GPCR families (Fig 4).

**Fig 4.**
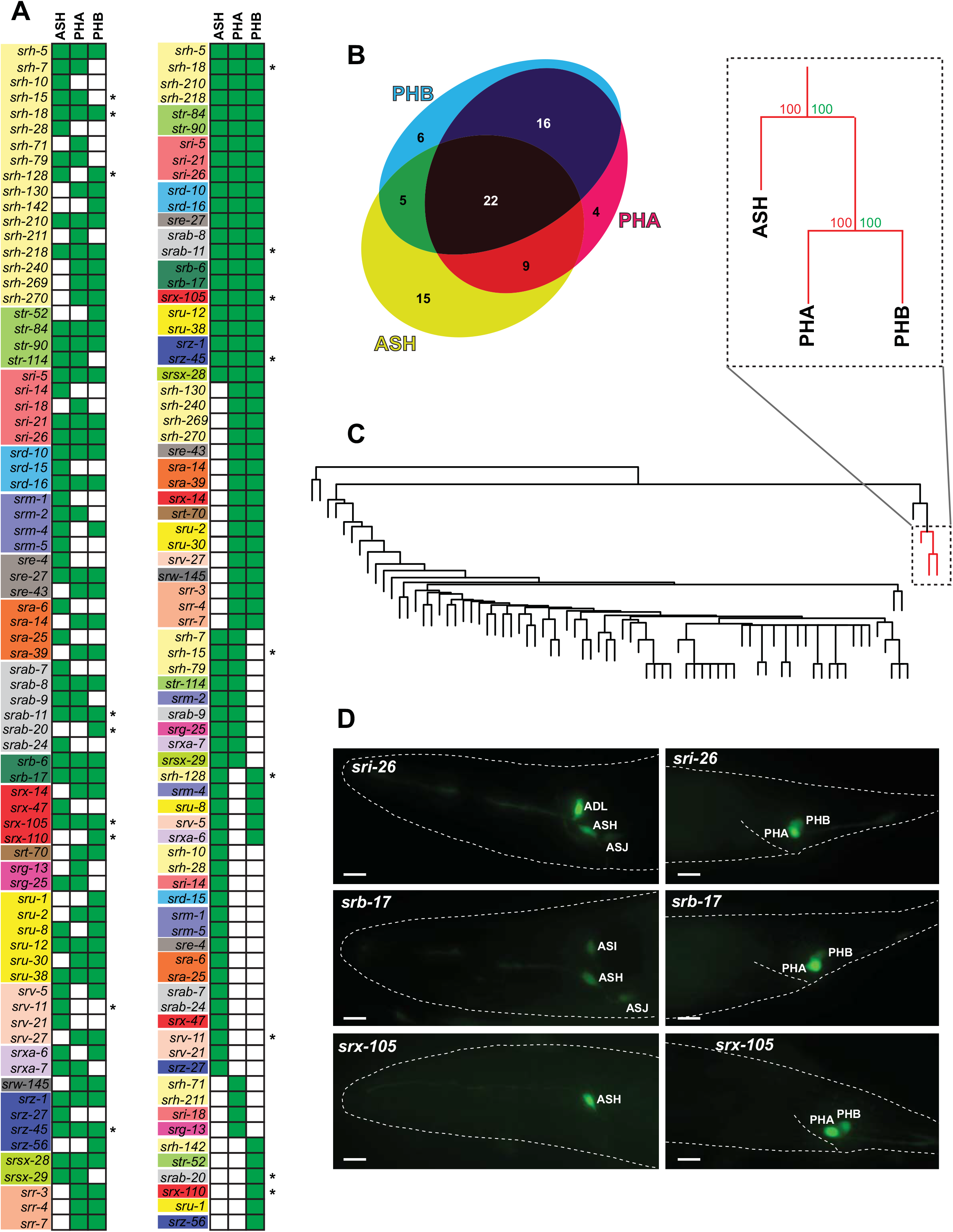
The only coexpression association of GPCR reporters is in nociceptive neurons. (**A,B**) Graphical representation of ASH, PHA and PHB co-expression. Green-filled square indicates expression. An asterisk denotes that the gene is exclusively expressed in the indicated neurons. Venn diagram was created with eulerAPE [57]. (**C**) Hierarchical clustering of neurons by GPCR reporter expression. Red lines show the well-supported ASH, PHA and PHB cluster (AU>95). AU: approximately unbiased p-value (percent), BP: Bootstrap Probability value (percent). (**D**) Examples of reporter gene expression profiles in ASH/PHA/PHB. Scale bars, 10μm.

### Left/right asymmetric expression of csGPCR reporters

One major motivation for undertaking the csGPCR reporter analysis was to identify more lateralized neuron pairs in the nervous system. In vertebrates, there is a striking dearth of molecular correlates for widespread functional lateralization of the brain. In *C. elegans* the chance discovery of left/right asymmetric sensory receptor expression has opened up new vistas on lateralization of the *C. elegans* nervous system [58]. Specifically, the lateralized expression of several csGPCRs in the AWC olfactory neuron pair [31] and guanylyl cyclase receptors in the gustatory ASE neuron pair [59] revealed a common theme of lateralization providing means of sensory discrimination [32, 60, 61]. Since lateralization provides an elegant, straight-forward means for sensory discrimination, we speculated that such lateralization may be wide-spread in the nervous system and therefore took particular care in examining whether csGPCR reporters that we analyzed are expressed in a left/right asymmetric manner.

We indeed identified eight csGPCR reporters with left/right asymmetric gene expression in an otherwise bilaterally symmetric neuron pair. However, this laterality was only observed in the context of the AWC sensory neuron pair, which was previously known to express several GPCRs in a left/right asymmetric manner [31, 62]. Using previously described sets of mutants, we found that the asymmetry of these GPCR reporters is controlled by the same calcium-dependent signaling pathway [33] that controls all other previously known asymmetric GPCRs in the AWC neurons (Fig 5). Of course, our limited analysis does not exclude the existence of left/right asymmetrically expressed GPCR genes in other neuron types, but it may not be as widespread as we initially hypothesized.

**Fig 5.**
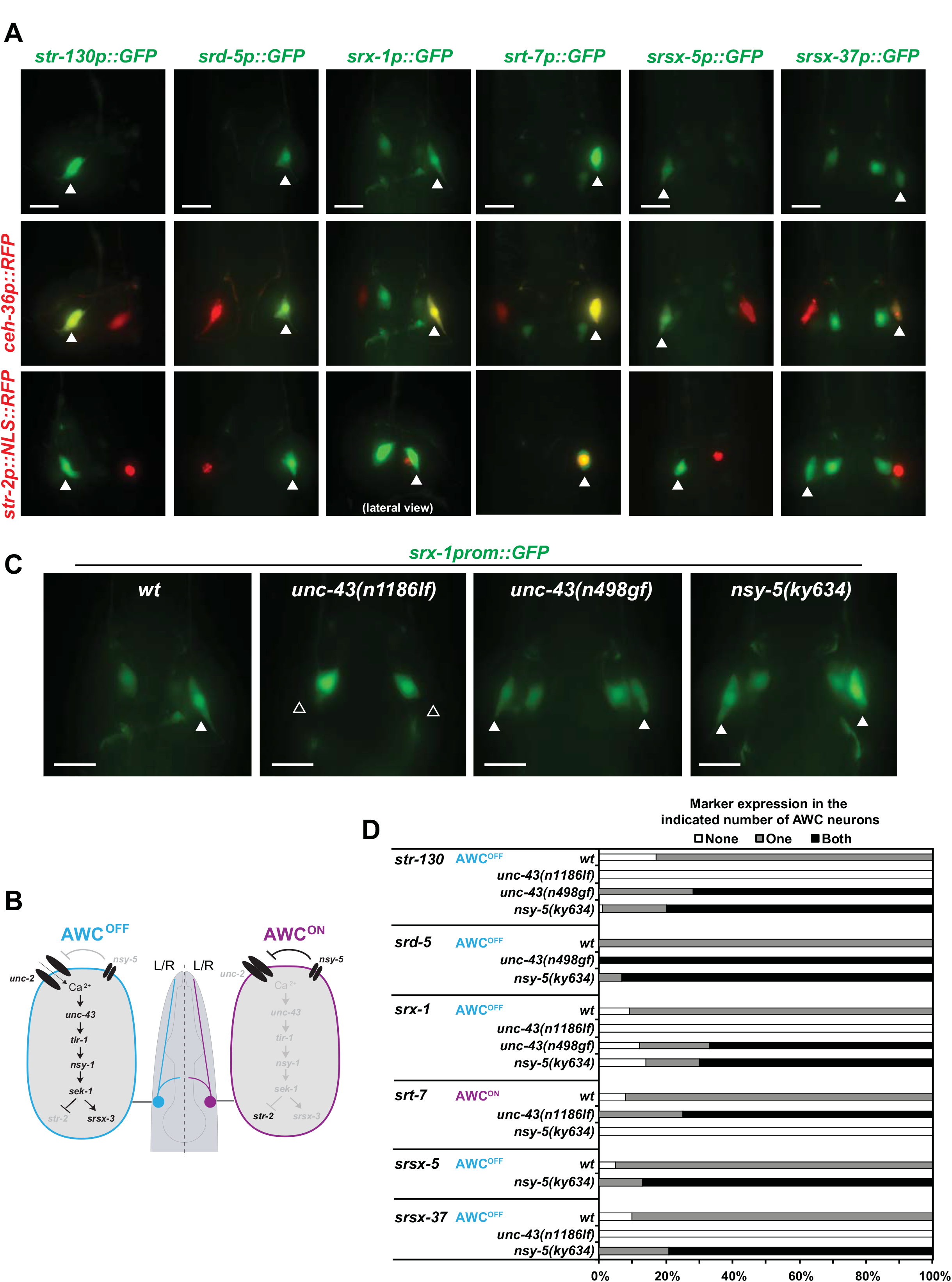
Lateralized GPCR reporter expression in the AWC neuron pair. (**A**) Asymmetrically expressed GPCRs, indicated with arrowheads (top row), were all expressed in AWC as assessed by colocalization with the *ceh-36p::RFP* reporter (middle row). *str-130, srd-5, srx-1, srsx-5* and *srsx-37* reporters were expressed in AWC^OFF^ while *srt-7* was expressed in AWC^ON^ as assessed with the *str-2p::NLS::RFP* reporter which is an AWC^ON^ marker (bottom row). All pictures are dorso-ventral views unless otherwise indicated. *srt-13* and *srr-9* reporters were also found to be asymmetrically expressed in AWC, however since these reporters were dim and not very robust no further analysis was done. Scale bars, 10μm. (**B**) AWC asymmetry. Previously known components of genetic pathways that control AWC asymmetries. Not all genes known to be involved are shown. Black and grey gene names indicate whether a gene is more active or more expressed (black) in one neuron compared with the other neuron. Scheme adapted from [63]. (**C**) Expression of the newly found AWC asymmetric GPCRs is regulated by previously described mechanisms. Representative pictures showing *srx-1* reporter expression (AWC^OFF^) in different mutants of the previously described AWC asymmetry pathway. As expected, in *unc-43(n1186lf)* mutants, *srx-1* reporter is expressed in none of the AWC neurons (2 AWC^ON^ phenotype) while in *unc-43(n498gf)* and *nsy-5(ky634)* mutants *srx-1* is expressed in both AWC neurons (2 AWC^OFF^ phenotype). Scale bars, 10μm. (**D**) Expression quantification of AWC asymmetric GPCR reporters in *unc-43(n1186lf), unc-43(n498gf)* and *nsy-5(ky634)* mutants. Animals were scored as young adults and show the expected 2 AWC^ON^ or 2AWC^OFF^ phenotype. Between 20 and 40 animals were scored per genotype.

### Sexually dimorphic expression of csGPCR reporters

Apart from brain lateralization, another domain of nervous system research displays a striking dearth of molecular markers. While the existence of sex-specific neurons is widely appreciated in the nervous system of most animals, including *C. elegans* [64], it is much less clear to what extent neurons that are shared by the two sexes of a given species display molecular differences. Recent anatomical work in *C. elegans* revealed intriguing synaptic wiring differences between sex-shared neurons in the two sexes [65], but even in *C. elegans* there is a dearth of dimorphic molecular markers of sex-shared neurons. Given the distinct priorities that males and hermaphrodites display toward food and mate searching [66] and given that a number of sex-shared sensory neurons are known to respond to different cues in a sex-specific manner [49, 67], we hypothesized that we may discover a multitude of sex-specifically expressed GPCRs. We indeed identified several GPCRs that are expressed in hermaphrodite-specific neurons (HSN, VC motor neurons) or in several male-specific neurons (Fig 6); however, we did not detect differences in GPCR expression in sex-shared neurons. We emphasize here, however, that we did not systematically analyze all 245 reporters that we analyzed in the hermaphrodite for differences in expression in the male, but rather focused on those GPCRs that show expression in 1-3 pairs of neurons in the hermaphrodites.

**Fig 6.**
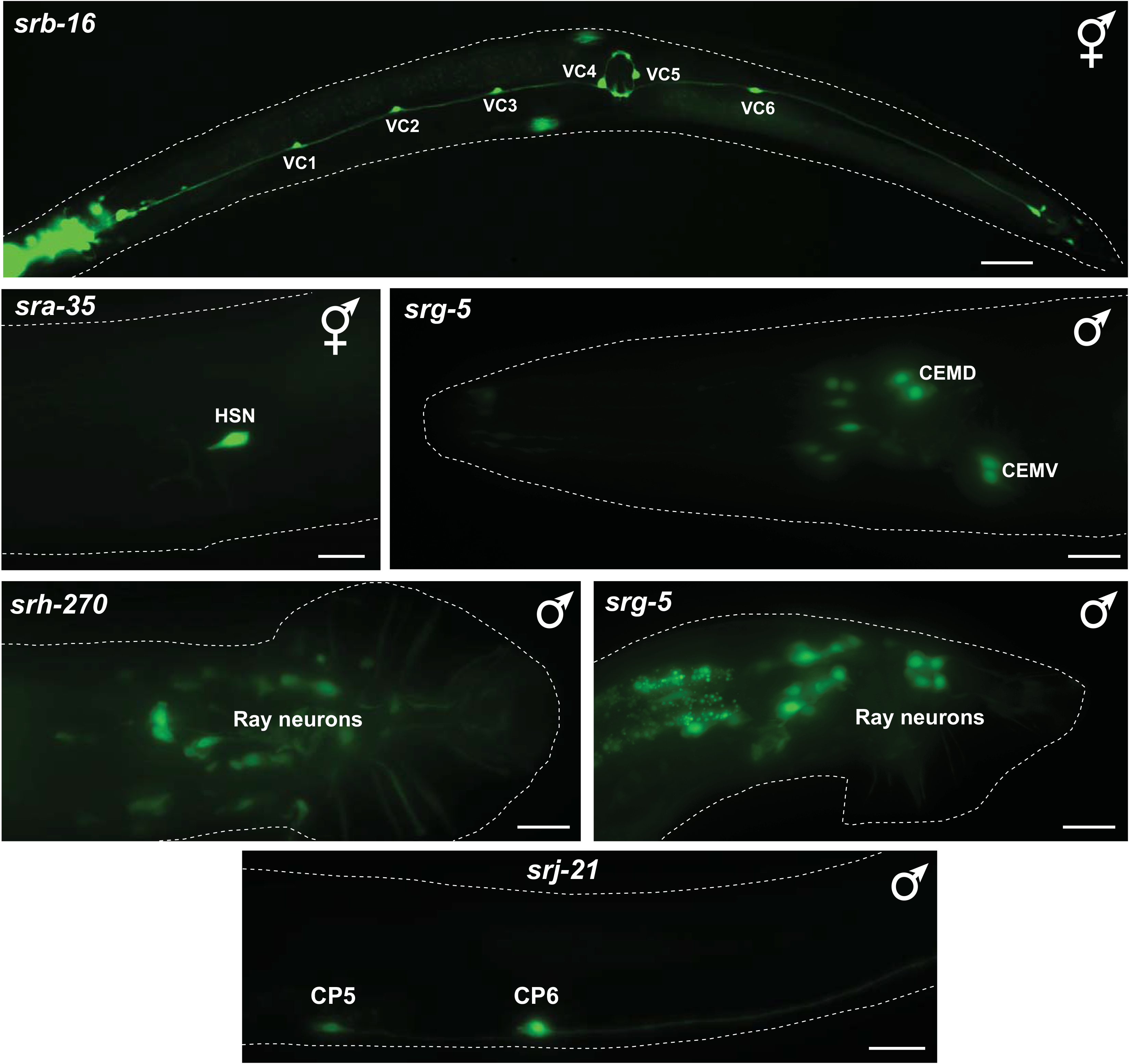
Expression of sex-specifically expressed GPCR reporters. Examples of GPCR reporters expressed in hermaphrodite-specific (VCs, HSN) or male-specific neurons (CEMs, CP5, CP6, Rays). All scale bars,10μm, except *srb-16* which is 30μm.

### csGPCR reporter expression outside the nervous system

Moving outside the nervous system, we found expression of individual GPCRs in essentially all tissue types (Fig 7 shows examples; summarized in Table 4). As we already mentioned above, the non-neuronal expression is often quite specific and there are only a few GPCRs that are expressed broadly in many different cell types (*e.g. srbc-58, srr-4*). Specific sites of non-neuronal expression include subsets of muscle cells, hypodermal cells, specialized epithelial cells, cells of the somatic gonad (distal tip cells), individual cells of the excretory system, glial cells and others (Fig 7, Table 4). There are no obvious, specific associations of non-neuronal expression with expression in a specific set of neuron types. Also, non-neuronally expressed GPCR receptors are not biased toward a single subfamily. GPCRs expressed in non-neuronal tissues that are exposed to the environment, e.g. epidermis, could be involved in sensing external cues but other non-neuronal cells will rather respond to internal signals. As a cautionary note, we can not presently exclude that nonneuronal expression may be the result of lack of repressor elements in the reporter constructs, but we note that in *C.elegans* there is presently little evidence for non-neuronal repressor mechanisms restricting gene expression to the nervous system (e.g. [68]).

**Fig 7.**
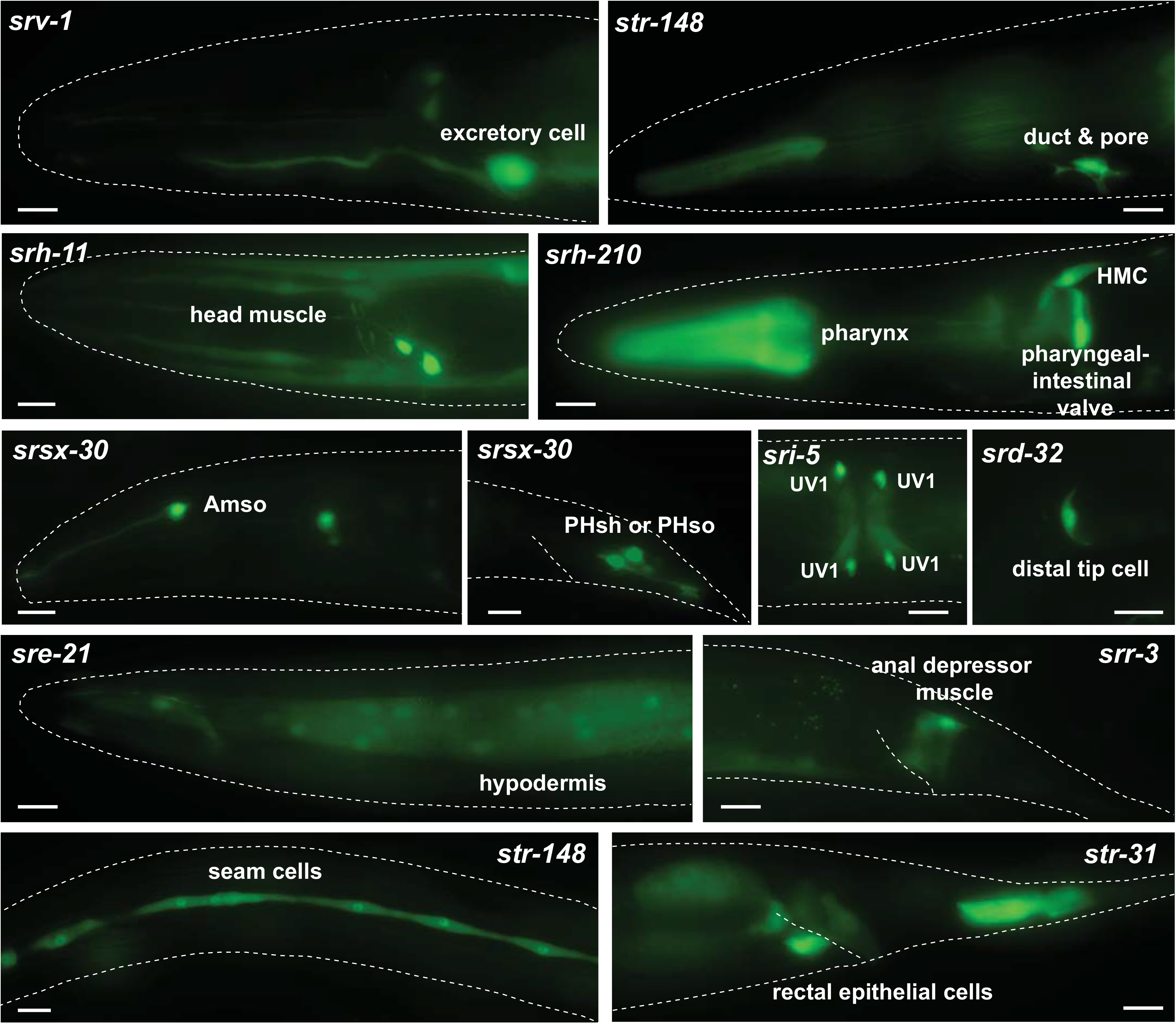
Expression of non-neuronal GPCR reporters. Examples of GPCR reporters expressed in different types of non-neuronal tissue. Scale bars, 10μm. See Table 4 for a complete summary of GPCR reporters expressed in non-neuronal tissues.

**Table 4:**
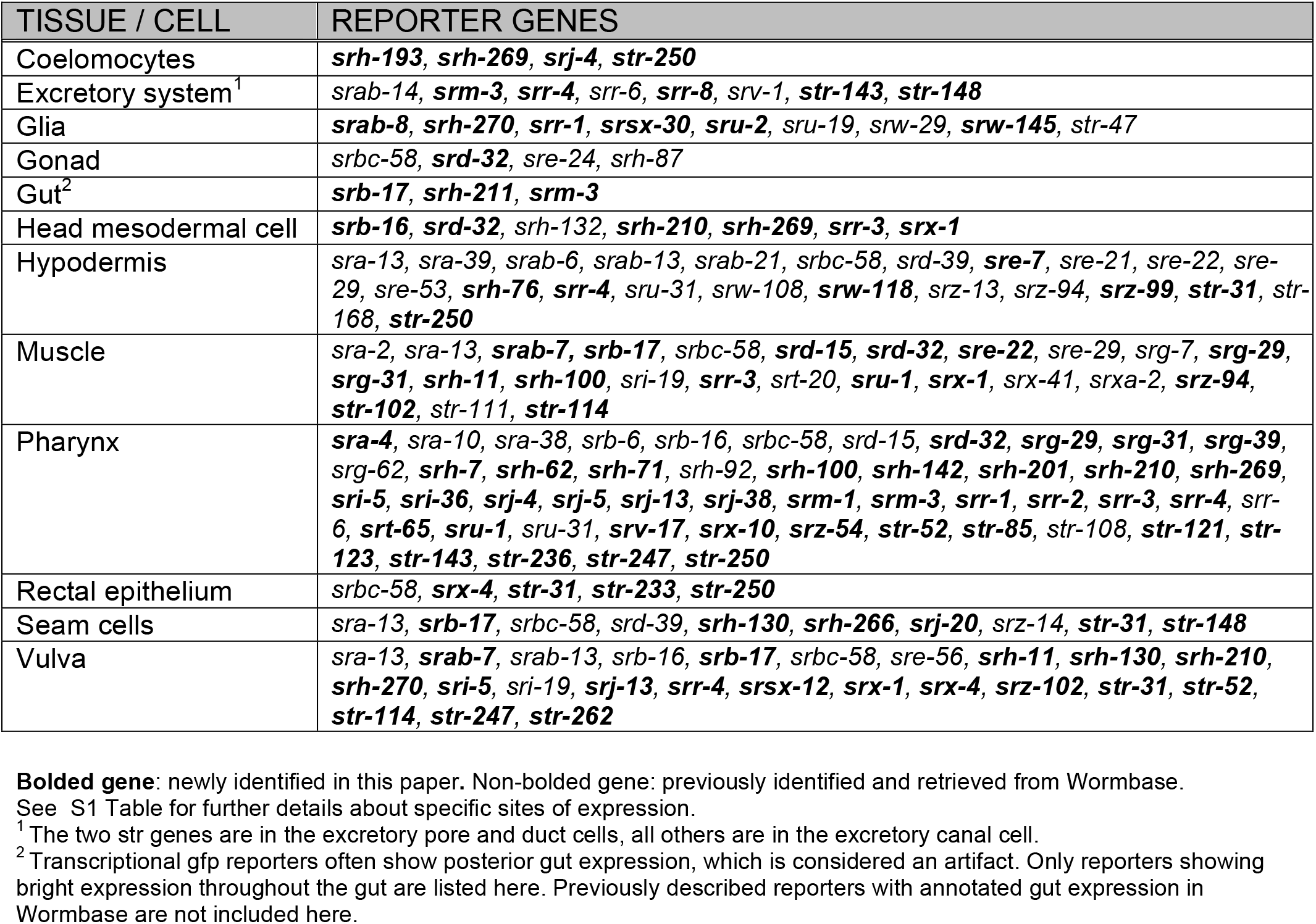
Non-neuronal sites of GPCR reporter expression

### Reporter gene analysis of entire csGPCR gene families

Do any of the patterns described above cluster with sequence similarity (*i.e*. family membership) of the receptors? As described above, specific features of csGPCR expression patterns do not correlate with family membership, but we wanted to pursue this issue further via a more comprehensive analysis of entire chemoreceptor gene families. As defined by sequence analysis [7], chemoreceptor gene families have very different sizes, ranging from a single gene per family (*srn* family) to 223 genes per family (*srh* family)(Table 2). We analyzed reporter gene expression patterns of all members of two small families to examine whether there are common themes in their expression patterns, their genomic location and *cis*-regulatory control regions. We also analyzed the expression of the one family, the *srn* family, which only has a single member, which is highly conserved in *Caenorhabditis* species, to assess whether it may show an unusual expression pattern. However, we find the *srn-1* reporter gene to be mainly expressed in amphid sensory neurons, like many other GPCRs (Fig 8).

**Fig 8.**
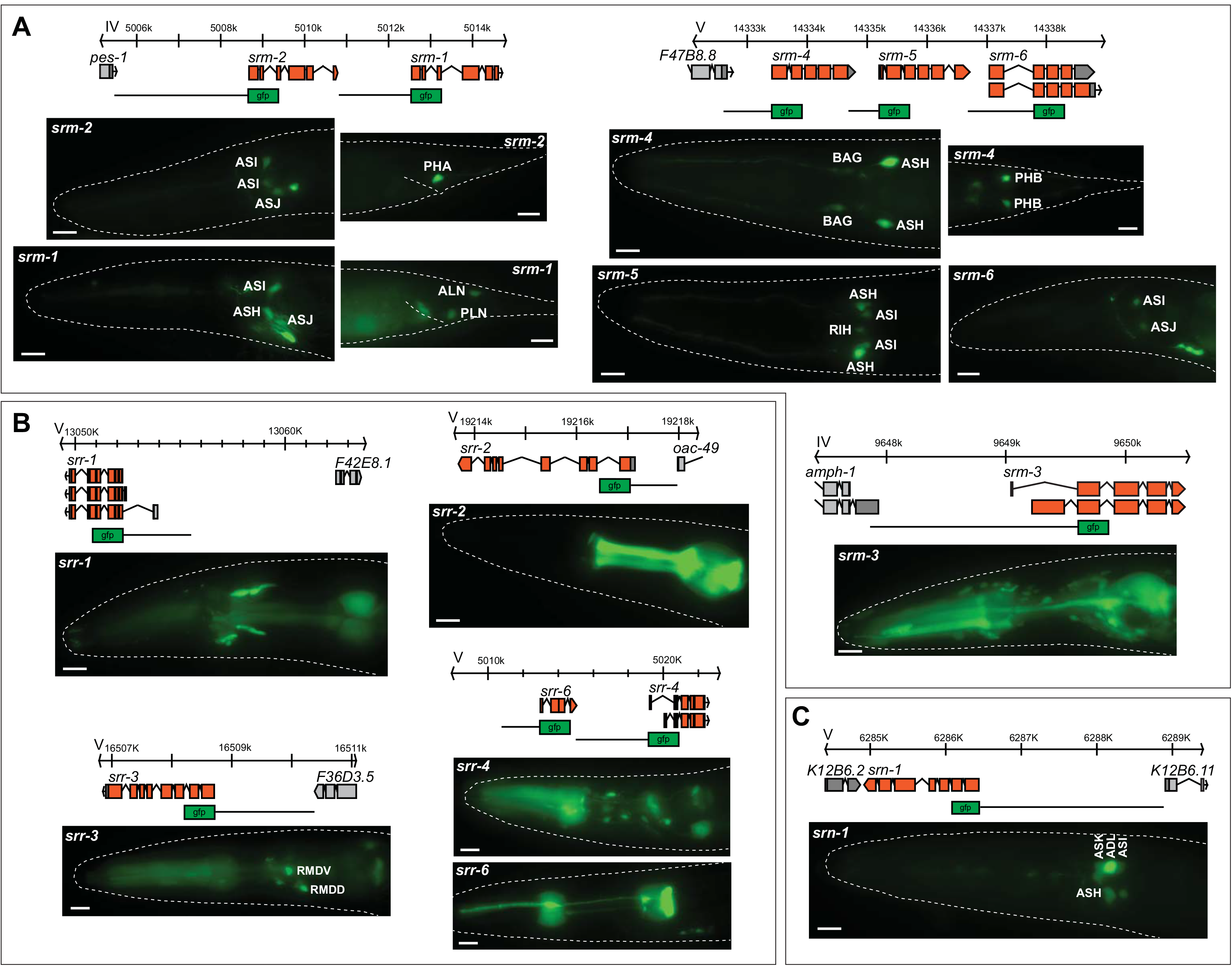
Reporter analysis of entire GPCR families. Genomic loci, reporter scheme and gfp expression images for the *srm* (**A**), *srr* (**B**) and *srn* (**C**) GPCR gene families. Only reporters expressed in the pharynx are shown for the *srr* family. Scale bars, 10μm.

The two small families for which we generated and analyzed reporter genes for all family members are the previously uncharacterized *srm* (6 members) and *srr* (9 members). Five out of the six *srm* family genes are syntenic to other family members (Fig 8). As these direct genomic adjacencies suggest local gene duplication, we could ask the question whether such local duplications also resulted in duplication of the 5’ *cis*-regulatory control regions and to what extent such duplicated *cis*-regulatory control regions retained similar expression profiles. We find that the adjacent *srm-1* and *srm-2* genes are expressed in a small set of mostly sensory neurons; some of these neurons are the same, others are different. The same theme applies to the adjacent *srm-4, srm-5* and *srm-6* genes. Their 5’ upstream regions direct expression to distinct, but partially overlapping sets of neurons.

The *srr* gene family is composed of 9 members. Reporter genes for all members displayed expression in diverse sets of neuron types with no common theme emerging. Outside the nervous system, it is notable that half of the family members are expressed in distinct cell types of the pharynx (Fig 8), suggesting a role for these genes in sensing food.

### Temporally regulated csGPCR reporter genes

We also sought to examine dynamic aspects of csGPCR expression. We focused on dynamics that relate to developmental timing and the response to harsh environmental conditions. To facilitate the identification of changes in expression, we focused our analysis on GPCRs that are robustly expressed in the adult in a small number of neurons (in most cases not more than 1-3 neuron pairs in the head and/or 1-2 neuron pairs in the tail). At the first larval stage we didn’t detect any differences in expression in 79 out of 82 examined reporters. Due to the limitations of multicopy array based fluorescent reporters, moderate intensity changes within a cell type might be difficult to notice and could have been missed. Three reporter genes, *srh-11, sru-48*, and *sra-28*, show striking differences in first larval *versus* adult stages: All three reporter genes show expression in the ASK neuron at the L1 stage, but not at the adult stage (Fig 9). Additionally, *srh-11* is expressed brightly in the ASI neuron at the L1 stage but dimly at the adult stage (Fig 9). Furthermore, dim expression of *srh-11* and *sra-28* reporter genes in the tail phasmid PHB and PHA neurons respectively, is only observed at the L1 stage but not at the adult stage (Fig 9).

**Fig 9.**
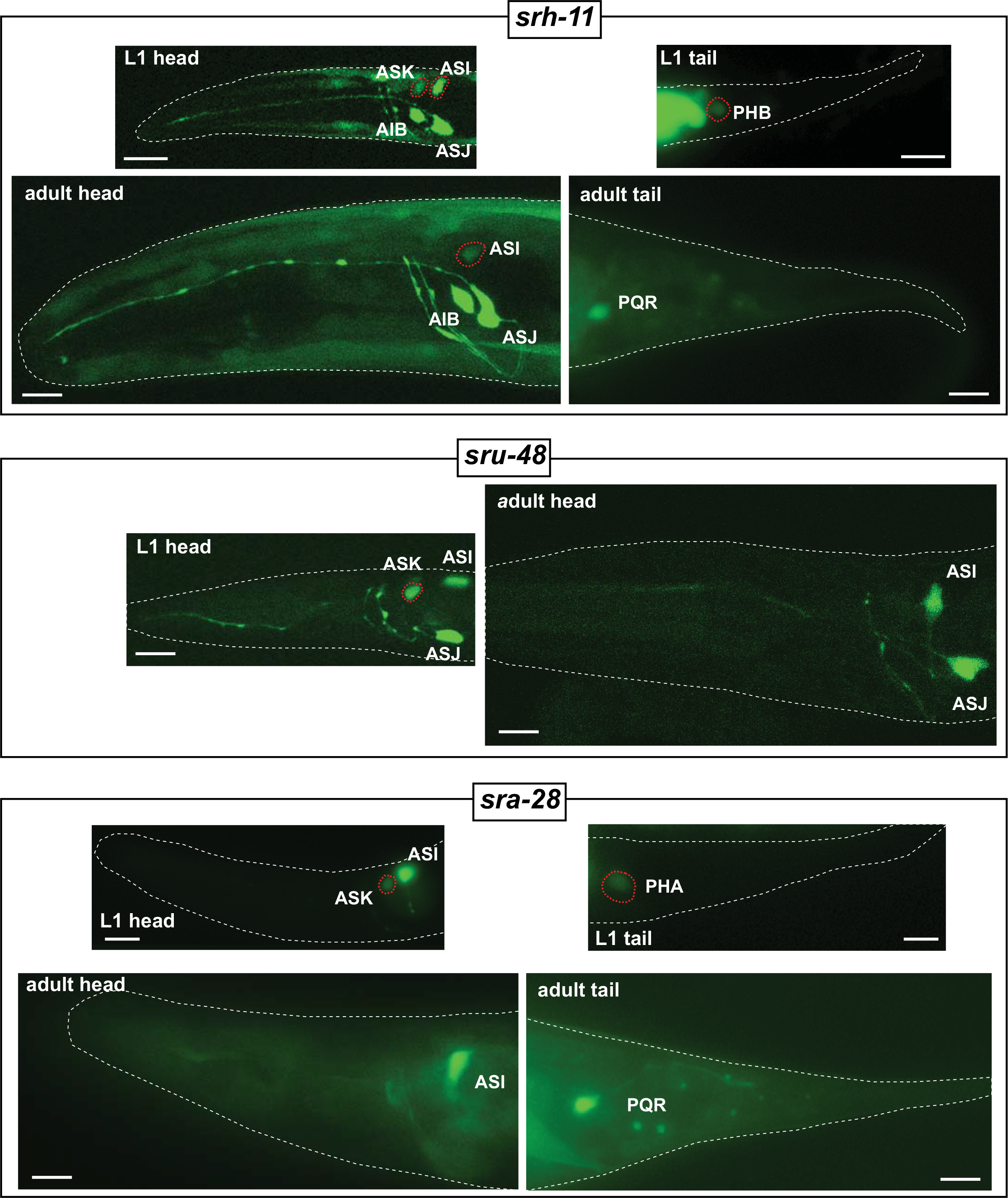
Temporal regulation of GPCR reporters. GFP images showing temporal expression changes (L1 versus young adult) of *srh-11, sru-48* and *sra-28* reporter genes. Neurons showing temporal changes in expression are outlined with red dotted lines. Scale bars, 10μm.

### GPCR reporter gene expression changes in dauers

We found that a substantial number of GPCR reporter genes were dynamically expressed when animals enter the dauer stage, an environmentally controlled diapause arrest stage that is accompanied by substantial cell, tissue and behavioral remodeling [69, 70]. Initially again focusing on reporters that are expressed in a restricted number of neurons under well-fed conditions, we found that 16 out of 46 examined reporters show a diverse set of changes in animals that were sent into the dauer stage via a standard starvation/crowding protocol (see Experimental Procedures). Many of the changes entail striking changes in the cellular specificity of GPCR reporter expression (Fig 10, Table 5). The vast majority of differences are observed in the nervous system, but some changes also occur outside the nervous system. Changes in GPCR reporter expression in the dauer stage have previously been described for two GPCR reporters [34](summarized with our novel patterns in Table 5), but the patterns we observe here are much broader and more complex. They can be summarized as follows:

**Fig 10.**
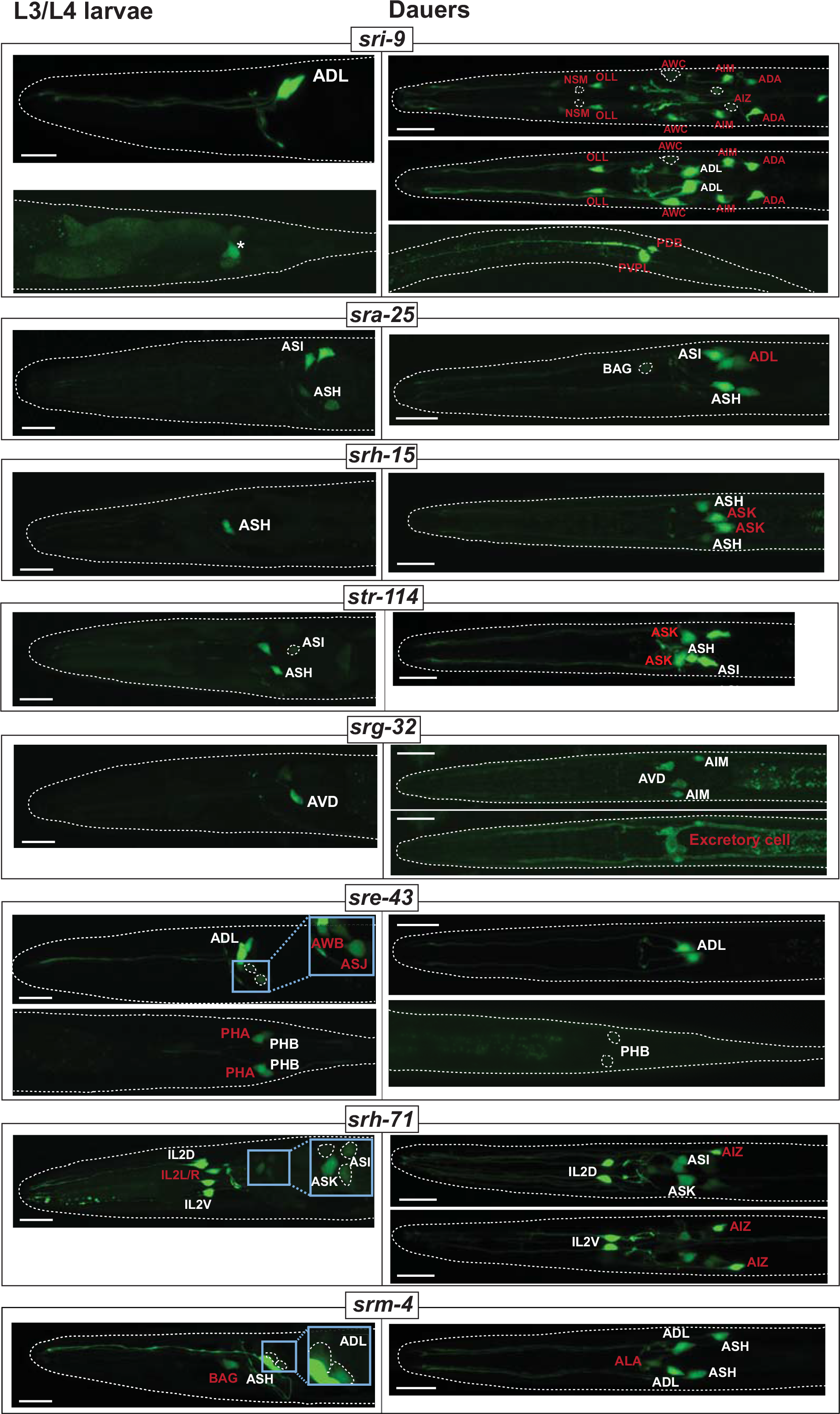
Examples of environment-induced changes in GPCR expression. Examples of GPCR reporters that change expression in dauer. Designations of neuron types that change expression are highlighted in red. Asterisk indicates posterior gut autofluorescence. Insets for *srh-71, sre-43* and *srm-4* show enlarged and overexposed images of cells that are too dim to be discernable in main panels. See Table 5 for a complete summary of GPCR expression changes in dauer. Scale bars, 10μm.

**Table 5:** Changes in GPCR reporter expression in starvation-induced dauers, within and outside the nervous system. Reporter gene expression patterns were analyzed in starvation-induced dauers. Previously reported GPCR reporter changes are listed in the two bottom rows of the Table [34]. For the *srh-71* reporter, we also observe non-robust expression in a non-phasmid pair whose identity we have not determined.

| Type of change | Reporter Gene | Reporter expression |  |  | Postdauer recovery |  |  |
| --- | --- | --- | --- | --- | --- | --- | --- |
|  |  | Constitutive expression in all stage (in dauer and non-dauer, fed L3 neurons and post-dauer) | Cells only show expression in dauers | Constitutively expressed in fed animals only, i.e. downregulated specifically in dauers in respective cell | Recovers | Dauer pattern is retained post-dauer | Entirely new post-dauer expression |
| dauer gains (type I) | *sri-9* | ADL | NSM (dim), OLL, AWC, AIM, AIZ, ADA, PDB, PVPL |  | NSM, OLL, AWC, AIM, AIZ, ADA turn off again | PVPL remains on, PDB occasionally on | none |
|  | *srh-15* | ASH, PHA | ASK |  | recovers to fed condition | none | none |
|  | *str-114* | ASH, ASI, PHA, head muscle | ASK, ASG |  | recovers to fed condition | none | none |
|  | *sra-25* | ASH, ASI, BAG (dim) | ADL, PHB |  | PHB turn off again | ADL remains on, BAG becomes bright and stable | ASJ (dim) |
|  | *str-84* | ASH, ASI, PHA, PHB, PVQ | Body wall muscle |  | recovers to fed condition | none | none |
|  | *srg-32* | AVD, AIM, PVQ | Excretory cell |  | recovers to fed condition | none | none |
| dauer losses (type II) | *sre-43* | ADL, PHB (dim in dauers) |  | AWB, ASJ (variable), PHA | AWB, PHA turn on again | ASJ become stable | ASH (dim) |
|  | *srh-279* | ADL |  | ADL down (but not off) in dauers | recovers to fed condition | none | none |
|  | *srz-67* | ADL |  | ADL down (but not off) in dauers | recovers to fed condition | none | none |
|  | *srx-12* |  |  | ADF, amphid sheath glia | recovers to fed condition | none | none |
|  | *sra-7* |  |  | ASK down (but not off) in dauers. F and U rectal epithelial cells off | recovers to fed condition | none | none |
| both gains and losses (type III) | *srm-4* | ASH, PHB, ADL (dim) | ADL (bright), ALA | BAG | recovers to fed condition | none | none |
|  | *srx-4* | ASK, ASI | ADL | B and Y rectal epithelial cells | recovers to fed condition | ADL expression partially remains | none |
|  | *srh-71* | ASK, ASG, ASI (dim), IL2D/V | ASI (brighter), AIZ, two ventral ganglion MN pairs | IL2L/R, PHA | PHA expression comes back to fed state | ASI, AIZ, two ventral ganglion MN pairs remain on, IL2L/R remains off | pharyngeal gland cells (ventral g1) |
|  | *srsx-29* | ADF | ASH | PHA | recovers to fed condition | none | none |
|  | *sru-12* | ASI, ASH, ASJ, OLL, PHB | PVQ | IL2, PHA | IL2, PHA turn on again | PVQ remains on | PLN |
| Peckol et al. 2001 | *srd-1* |  |  | ASI | recovers to fed condition | none | none |
|  | *str-2* | ASI (dim) | ASI (brighter) | AWC | recovers to fed condition | none | none |

(1) In most cases, there is stable and unchanged expression in several neuron classes in dauer and non-dauers, but upon dauer entry, expression is either turned on in additional neuron classes (“type I” regulation) or becomes undetectable in subsets of specific neuron classes (“type II” regulation) (Table 5; Fig 10). There are also combinations of both changes (type III regulation): In one particularly striking example, the *srh-71* reporter is expressed in some sensory neurons in both dauer and non-dauers, but undergoes a striking respecification in dauers. Reporter expression becomes undetectable in the lateral IL2, PHA and an additional pair of tail neurons in dauer, and instead is turned on in the AIZ and ASG neurons (and increases expression levels in ASI). This hints toward the re-routing of internal sensory information.

(2) In a number of cases reporter expression is strongly downregulated, becoming undetectable in all neurons in which the reporter is expressed (Table 5; Fig 10).

(3) The changes outside the nervous system concern three tissue types, muscle, the excretory cell and epithelial blast cells (Fig 10). In two cases, expression of a specific csGPCR reporter is turned on in the dauer stage, while in another case expression becomes undetectable. These findings indicate that these tissue types now became receptive to signals in a dauer-specific manner, an unanticipated finding.

(4) The most recurrent set of changes in the expression of distinct reporters concern nociceptive neurons, namely the ASH, ADL and phasmid tail neurons. Of particular note is the PHA phasmid neuron, which shows the most consistent pattern of changes: 4 csGPCRs are turned off or strongly downregulated specifically in the dauer stage.

(5) The most unusual novel expression pattern observed in dauer stage animals concerns the PVP tail interneuron pair. We found that in dauers, expression of the *sri-9* reporter is turned on in a left/right asymmetric manner, only in the PVPL neuron. The cellular identity of *sri-9* expression (as well as other expression changes) was corroborated by examining overlap of GPCR *gfp*-based reporters with *rfp*-based landmark strain (see Experimental Procedures).

### Some csGPCRs serve as molecular markers of life history

Do reporter expression changes observed in dauers recover upon re-feeding to the pattern observed in control fed animals? Examining csGPCR reporter expression in well-fed adult animals that had passed through the dauer arrest stage during larval development, we found that the expression of 11 of the 18 reporters, which showed dauer-specific gene expression changes, recovers to that of the fed state, i.e. in these 11 cases, expression in the adult is independent on whether the animals had passed through the dauer stage or not.

For 7 csGPCR reporters we discovered intriguing, cell-type specific alterations in animals which have passed through the dauer stage (Fig 11, Table 5). We observed two types of changes:

**Fig 11.**
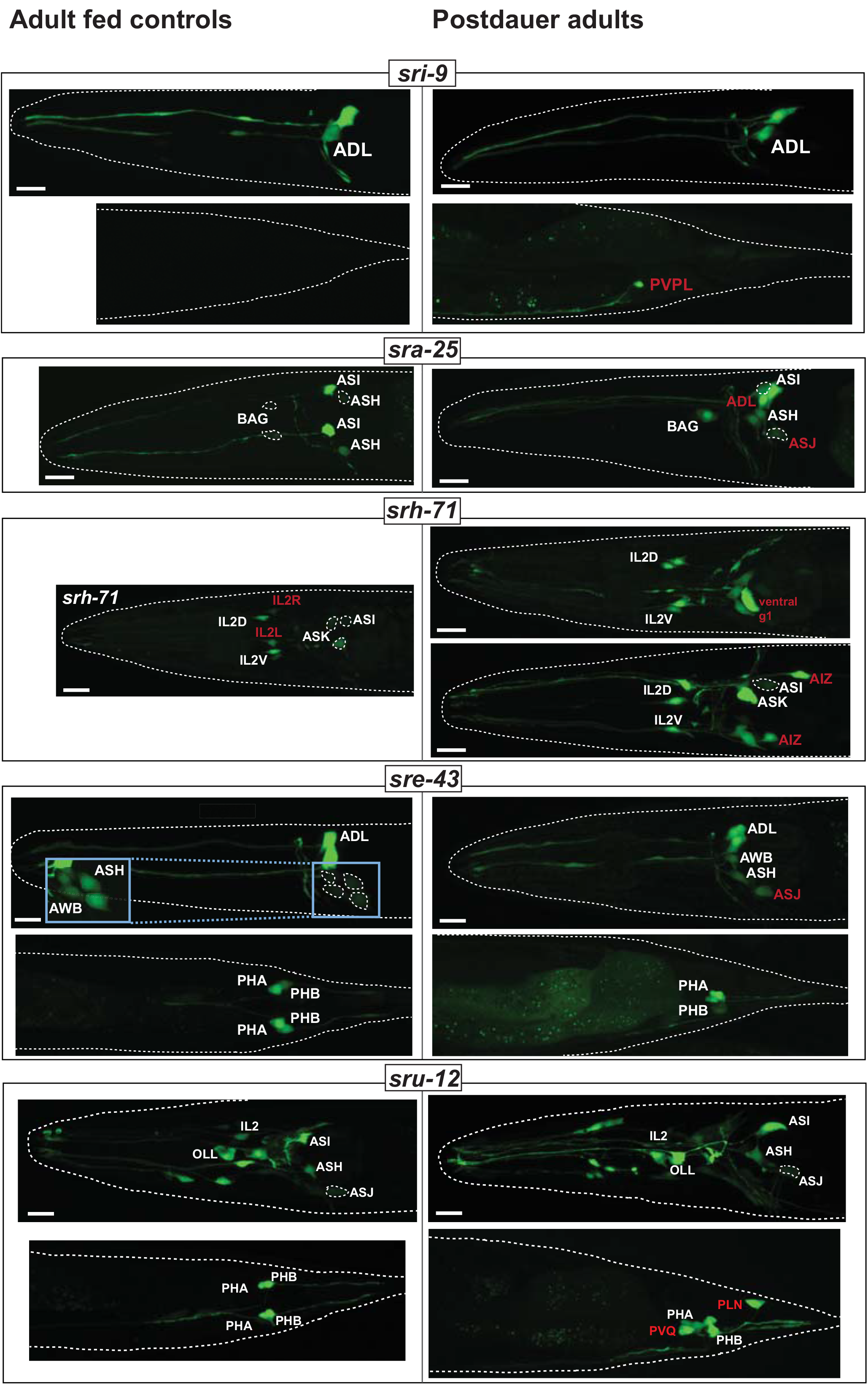
GPCR expression patterns as life history traits. Comparison of GPCR expression in 1-day old adult animals that either did pass through the dauer state (right panels) or did not (age-matched fed controls; left panels). Postdauer animals were in the dauer stage for 5-7 days. Designations of neuron types that retain dauer-specific expression or acquire postdauer-specific expression are highlighted in red. Inset for *sre-43* shows enlarged and overexposed images of cells that are too dim to be clearly discernable in the main panel. See Table 5 for a complete summary of GPCR expression changes in postdauer. Scale bars, 10μm.

(1) For 4 reporters (for *sri-9, sra-25, srh-71* and *sru-12*), we observed that expression which was induced in specific neuron types exclusively in dauers, retained this expression in postdauer animals: Dauer-induced expression of the *sri-9* reporter in PVPL, of the *sra-25* reporter in ADL, of the *sru-12* reporter in PVQ as well as of the *srh-71* reporter in AIZ and two ventral ganglion head motor neuron pairs is retained in post-dauer adults. In contrast, dauer-induced loss of *srh-71* expression in the lateral IL2 pair does not recover.

(2) In 4 cases (*sre-43, srh-71, sru-12, sra-25* reporters) we observed induction of expression in additional cell types exclusively in postdauer animals. *sru-12* reporter expression is specifically induced in the PLN neurons of postdauer animals, *sre-43* expression in dimly observed in the ASH neurons of postdauer animals, *sra-25* expression is dimly observed in the ASJ neurons in postdauer animals and *srh-71* reporter expression was induced in a non-neuronal pair, pharyngeal gland cells, in post-dauers.

In addition to these two categories, we found two instances, in which a sporadic and weak expression observed in animals that have not passed through the dauer stage will become highly penetrant and stable if they have passed through the dauer state (*sra-25* in BAG neurons, *sre-43* in ASJ, *srh-71* in ASI).

Note that all of the reporters for which we observe changes in postdauer recovery do recover their “fed patterns” in other neuron classes (these could be considered as internal controls that argue against the changes in expression being a reporter gene artefact). Taken together, adult animals show neuron-type specific differences in the expression of GPCR reporters depending on whether they have passed through periods of distress. GPCR reporters therefore serve as reporters of life history traits.

### L1 starvation recapitulates some but not all csGPCR reporter changes

We tested five of the 16 csGPCR reporters that displayed changes in the dauer stage for whether their expression also changes in another starvation-induced arrest stage, the starvation-induced L1 arrest stage. Comparing expression in 2 day-starved L1 (egg prep into M9 medium) to fed L1, we find that two reporters (*str-114* and *sra-25*) show the same changes as observed in dauer animals (Fig 12). In contrast, two reporters (*str-84* and *srg-32*) that change their expression in dauers, do not show changes in starved L1 vs. fed L1 (Fig 12). One reporter, *srh-15*, in addition to dauer-specific expression in ASK, is also expressed in ASI in starved L1. Hence, the response of GPCR expression to arrest conditions is diverse.

**Fig 12.**
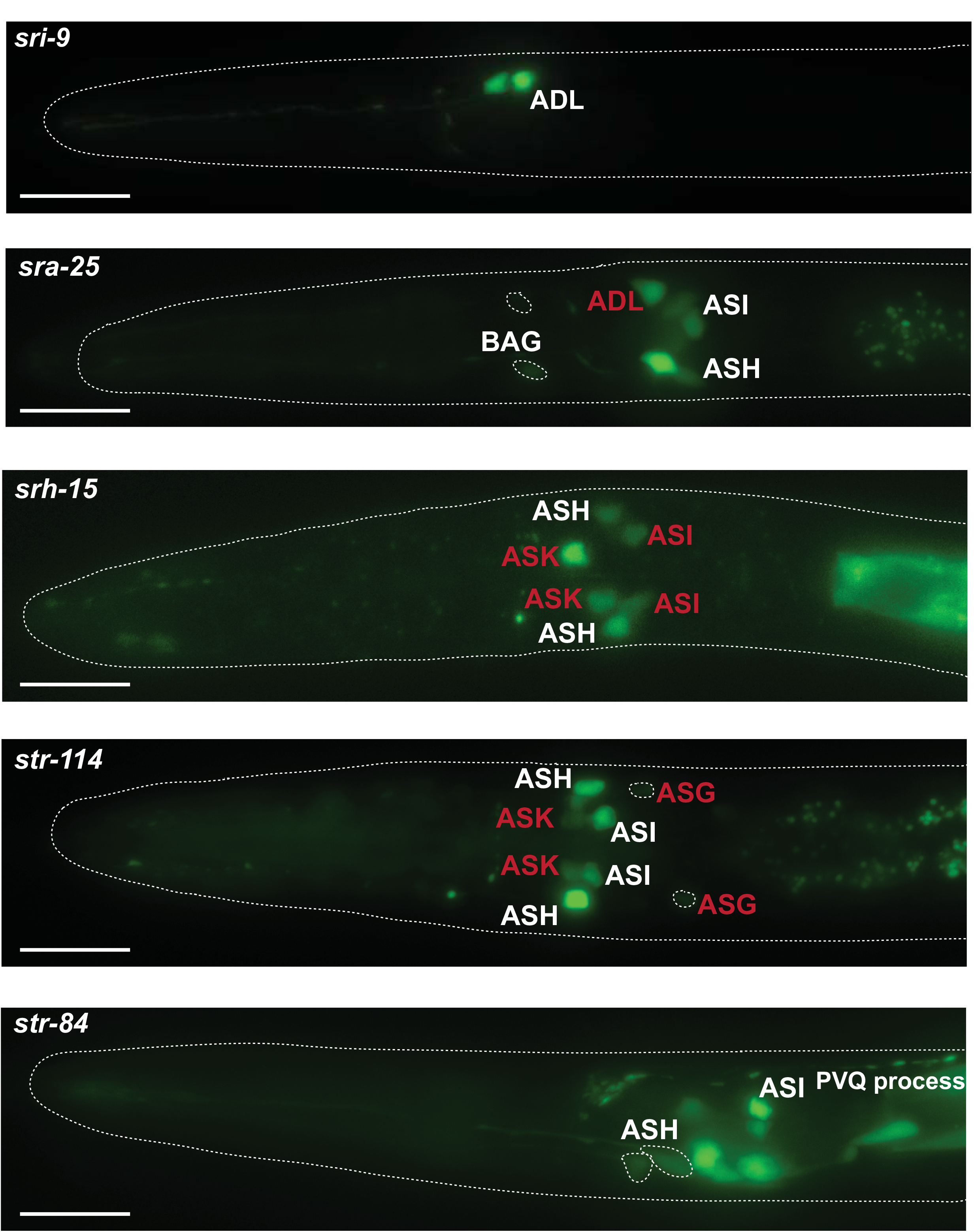
GPCR reporter expression in starved L1 animals. Examples of GPCR reporter expression in starved L1 worms. Eggs isolated by bleach treatment were allowed to hatch and were kept in M9 for 48 h. Designations of neuron types that change expression compared to fed L1 worms are highlighted in red. Scale bars, 10μm.

## DISCUSSION

Together with previously published analyses, there are now reporter transgenes that monitor the expression of 373 of the ~1,300 chemosensory GPCR genes encoded in the *C. elegans* genome. One intrinsic limitation of reporter genes is that they do not necessarily capture the full complement of *cis*-regulatory control elements of a gene. However, given the compact nature of csGPCR loci, the inclusion of all 5’ regions in most reporters and the small size of introns, the number of inaccuracies may be quite limited. Irrespective of whether the reporters are a reflection of the complete expression of a csGPCR, they nevertheless function as highly valuable molecular markers of cellular identity and plasticity. Meaning, reporter gene analysis decodes *cis*-regulatory information and provides read-out of regulatory states of specific cell types. The key conclusions of the expression patterns inferred from the reporter genes are as follows:

### Restricted expression

Most csGPCRs show a very restricted expression in few cell types. Many GPCRs are expressed in single neuron classes. Those csGPCRs that express in multiple neuron types do not display a coherent set of coexpressing neurons, with one notable exception: the nociceptive ASH, PHA and PHB neurons display similar (but not identical) sets of csGPCR expression.

### csGPCR coexpression within a neuron class

Some neurons display a remarkably large number of GPCRs. The ASI neuron displays the most csGPCR genes at 99, followed by many distinct types of nociceptive neurons. While csGPCRs have been found for all but two sensory neurons (URY and ADE), there is a striking disparity in the number of csGPCRs coexpressed in sensory neuron types. Amphid sensory neurons clearly coexpress the largest number of GPCRs, while other sensory neurons express many fewer csGPCRs. The nociceptive ADL stands out in the list of amphid neurons, as it is the neuron expressing the most single-neuron specific GPCR reporters.

### Expression in sensory and non-sensory neuron classes

While expression of csGPCRs clearly predominates in sensory neurons, they are also expressed in inter- and motorneurons and in a diverse set of non-neuronal cells. In most cases, each GPCR is restrictively expressed, suggesting that many different cell types in an organism show very distinct and cell-type specific responses, likely to internal signals. The similarity of one GPCR family, the *srw* family, to peptide receptors of other animal species provides hints to the nature of these ligands [11, 29]. The expression of many members of the *srr* family in pharyngeal tissues suggests another source of ligands; perhaps these receptors respond to cues from ingested bacteria. In vertebrates, chemosensory GPCRs are now also becoming increasingly appreciated as being expressed in non-neuronal cells [6].

### Polymodality of sensory neurons

csGPCRs were detected in sensory neurons that are known to express distinct types of sensory receptors and engage in non-chemosensory behavior, *e.g*., in gas-sensing neurons or different types of mechanosensory neurons. The expression of csGPCRs in these neuron classes may hint toward these neurons perceiving different sensory inputs, *i.e*., they are likely polymodal. However, as discussed above, csGPCRs expressed in these neurons may not be involved in detecting external sensory cues, but measuring internal states.

### Absence of gene family themes

The absence of any overarching expression theme within gene families is striking. We did not observe that the expression of family members clusters in specific neuron types or share any other specific expression features. Specifically: (a) left/right asymmetrically expressed csGPCRs in the AWC neurons do not fall into the same family; (b) csGPCR reporters that are differentially regulated in larval stages or in the dauer stage do not come from a single family; (c) GPCRs that share specific expression pattern themes (*e.g*., coexpression in the nociceptive ASH, PHA and PHB neurons) do not derive from specific families; (d) non-sensory neuron-expressed or non-neuronal expressed GPCRs do not fall into a specific family. The only glimpse of perhaps some common function is observed in the small *srr* family (9 genes), half of which appear to be expressed in nonneuronal pharyngeal tissue. An important note of caution is that these conclusions are based only on reporter genes. However, the substantial sample size on which these conclusions are based lends some credence to these conclusions.

### Combinatorial complexity

GPCRs generally act as homo- or heterodimers [71], thereby hugely increasing the amount of distinct sensory receptor complexes expressed in a cell. This combinatorial activity also makes prediction of function of a given GPCR very difficult in that a GPCR may have one function expressed in one cell (in combination with another GPCR), while it may have a very different function in another cell (in combination with yet another GPCR).

### Left/right asymmetric csGPCR expression patterns

While we recovered novel csGPCR genes expressed in a left/right asymmetric manner in the AWC neuron pair, we were surprised to find no other obvious left/right asymmetries in other sensory neuron pairs. Of course such asymmetries may still be found with currently not analyzed GPCR genes, but the number of AWC asymmetries recovered suggest that AWC neurons may be exceptional in their extent of lateralization.

The only other asymmetry that we found revealed itself not in a sensory, but an interneuron and only in an non-anticipated context. The *sri-9* reporter transgene becomes induced in dauer animals in PVPL, but not PVPR, and PVPL expression is retained in postdauer animals. Molecular asymmetries in PVP neurons have not previously been reported but can perhaps be inferred by the fact that PVPL and PVPR are innervated in a left/right asymmetric manner by unilateral neurons. Specifically, PVPL, but not PVPR, is innervated by the unilateral DVB neuron. Perhaps *sri-9* may play a role in this synaptic signaling context, but why this should be dauer-specific is unclear.

### Plasticity of csGPCR expression

One notable feature of our analysis was the extent of plasticity that csGPCR reporters show in the context of the dauer stage. Dauer animals are thought to remodel most tissue types and significantly alter behavioral patterns. Changes in csGPCR expression and hence changes in the external and internal signal perception fit very well into the mold of organismal plasticity and illustrate the plasticity of many different tissue types (note, for example, the changes in csGPCR expression in muscle). We find it particularly intriguing that several csGPCRs represent markers of life history. Some of the changes in GPCR reporter gene expression in dauers is retained in postdauer animals and some csGPCR reporters turn on only in postdauer animals. Animal-wide expression transcriptomic analysis has previously identified large cohorts of transcripts that, like our csGPCR reporters, serve as marker of dauer life history, i.e. transcript changse in dauers and these transcript changes persisting in post-dauer animals [72]. However, due to the whole animal nature of this analysis, this previous study lacked cellular resolution. Our findings add a novel spatial component to these previous findings, since we find the life history traits to be strikingly neuron-type specific. The expression of the TRP channel gene *osm-9* has also previously been shown to be modulated during dauer and postdauer stages in a neuron class-specific manner; in this case, *osm-9* expression is downregulate in the ADL (but not AWA) chemosensory neurons and the repression is retained post-dauer, using RNAi-and chromatin-based mechanisms [73]. In all except one case that we report here, we observe the opposite post-dauer effect; reporters that are turned on in dauers, persist in non-dauers. The mechanistic basis of this may hence be distinct from the *osm-9* case.

It is important to remember here that the life history trait observations are based on transcriptional reporter genes which, on the one hand, may not accurately reflect expression of the endogenous locus, but, on the other hand, clearly provide a definitive molecular “read out” of changes in the “regulatory state” of specific neuron types, depending on whether they have passed through the dauer stage or not. Moreover our transcriptional reporter also argue that the life history regulatory phenomenon must be transcriptional in nature. These GPCR reporters will therefore provide excellent starting points to analyze the molecular mechanisms controlling this plasticity.

### Future uses of the csGPCR expression map

The csGPCR reporter atlas can be put to a number of future uses. The sites of expression of specific csGPCRs point to potential functions of the csGCPRs, guiding the future analysis of csGPCR knockout strains – many available by knockout consortia or easily constructable by CRISPR/Cas9 technology. For example, csGPCRs expressed in the polymodal nociceptive ASH, ADL and phasmid neurons may be mediating the response to a number of distinct sensory cues processed by these neurons [56, 74].

csGPCR expression patterns point to perhaps unexpected cellular sites of internal signal perception that warrant further investigation. For example, the excretory canal cell expresses at least six csGPCRs reporters (considering that we only examined reporters for ~20% of GPCR loci, this number may increase several fold). The relevance of this expression could be tested through the excretory cell-specific expression of dominant negative versions of G-protein downstream signaling components. Similarly, the cellular dynamics in csGPCR expression patterns point to specific cells undergoing changes that warrant future characterization. For example, the induction or suppression of csGPCR reporter expression during the dauer stage in specific sensory and interneurons that were not previously associated with dauer-specific functions may warrant a closer examination of other molecular and functional changes of these neurons during the dauer stage.

Because csGPCR reporter fusions also link precisely delineated sequences (used for reporter construction) to specific cellular sites of gene expression, patterns of coexpression of GPCRs can be used to extract *cis*-regulatory information, which in turn may point to *trans*-acting factors involved in controlling GPCR gene expression. A proof of principle for this type of analysis has already been conducted, pointing to a critical function of, for example, a bHLH factor in controlling csGPCR expression in the ADL nociceptive neuron [28] and with now substantially more expression information available can be further extended to additional cell types.

Lastly, GFP reporter transgenes have generally served as invaluable starting points for genetic mutant screens in which the genetic control of specific biological processes can be investigated. The GPCR reporter collection provides a multitude of entry points. For example, the postdauer expression of multiple reporter genes can be used to screen for mutants in which these life history traits fail to be properly expressed. GFP reporter genes have also served as invaluable cellular identity markers and here again the csGPCR reporter collection can be used to assess how the identity of specific cell types is genetically controlled.

## ACKNOWLEDGEMENTS

We thank Q. Chen for generating transgenic strains, the Vancouver gene expression consortium for generating strains and the CGC for providing these strains to us. We thank Neda Masoudi and Sarah Finkelstein for communicating unpublished results.

---

**S1 Fig.**
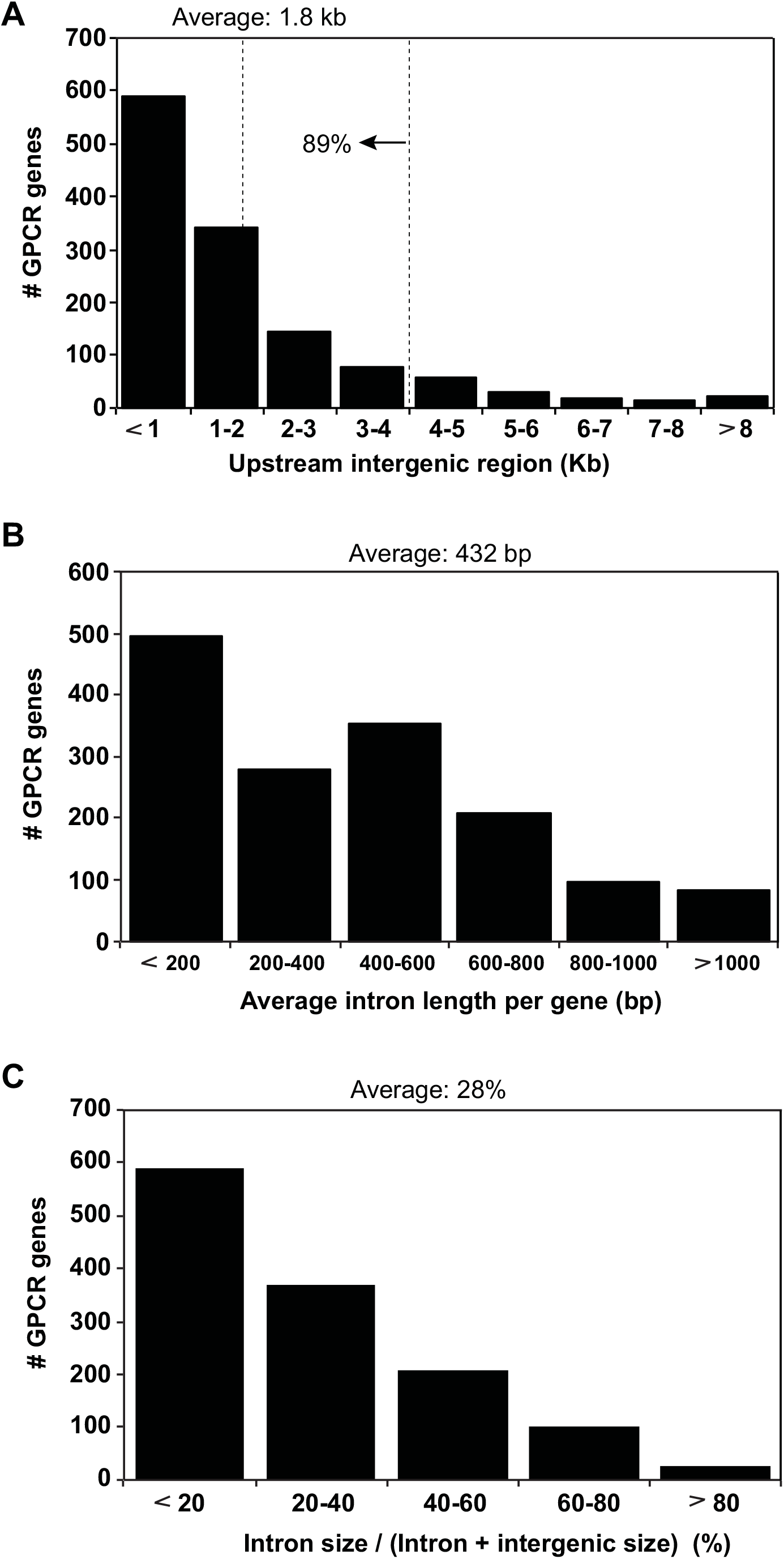
csGPCR gene locus analysis. (**A**) Histogram of upstream intergenic region distances of all *C. elegans* cs GPCR genes. The average size of the 5’ intergenic region (= distance to next gene) is 1.8kb. 89% of all loci have a 5’ intergenic region of smaller than 4kb. (**B**) Histogram of average combined intron length (bp) per GPCR gene. (**C**) The intergenic region of the majority of GPCR is substantially larger than the combined intronic region.

**Table S1:**
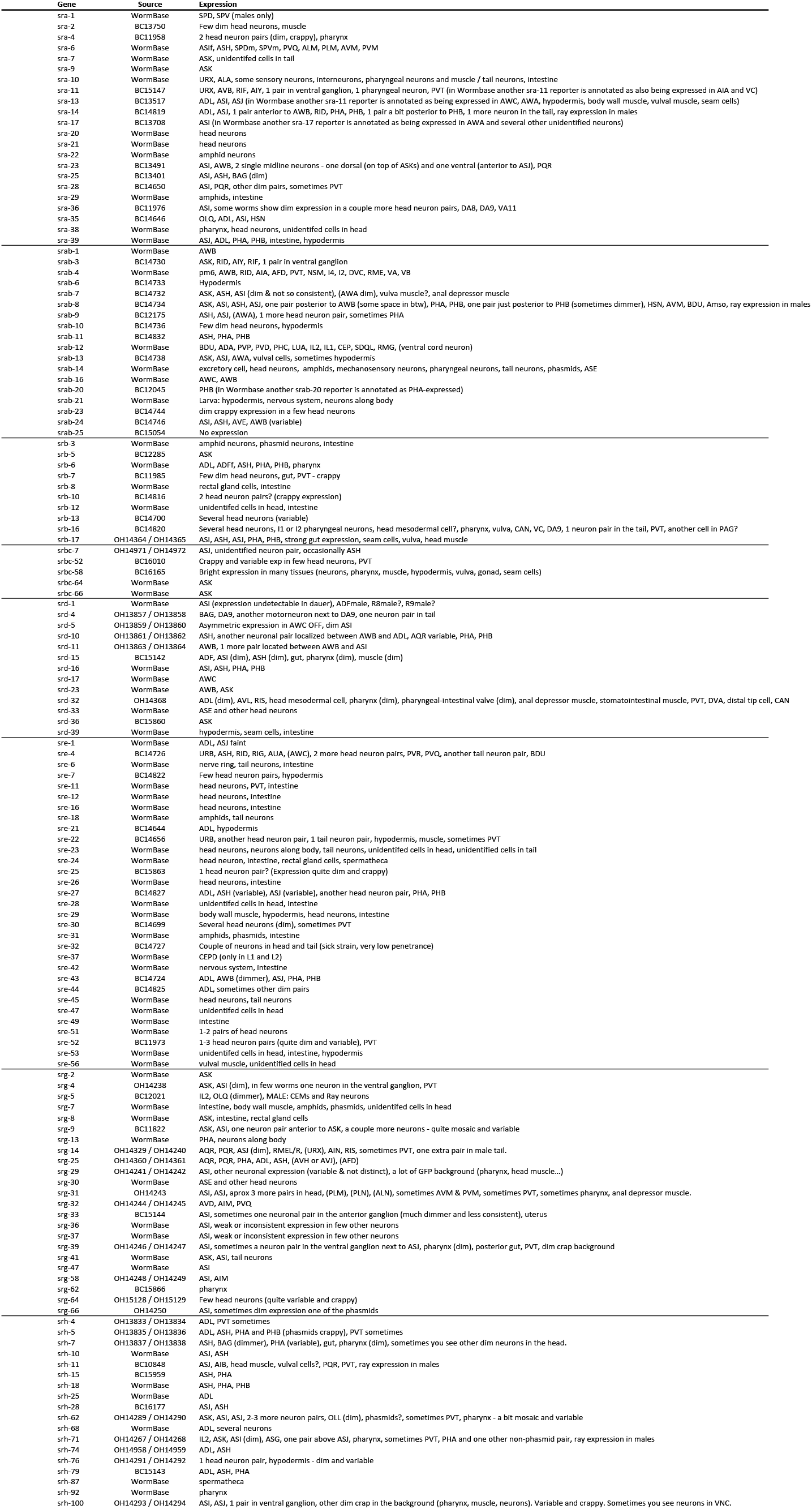
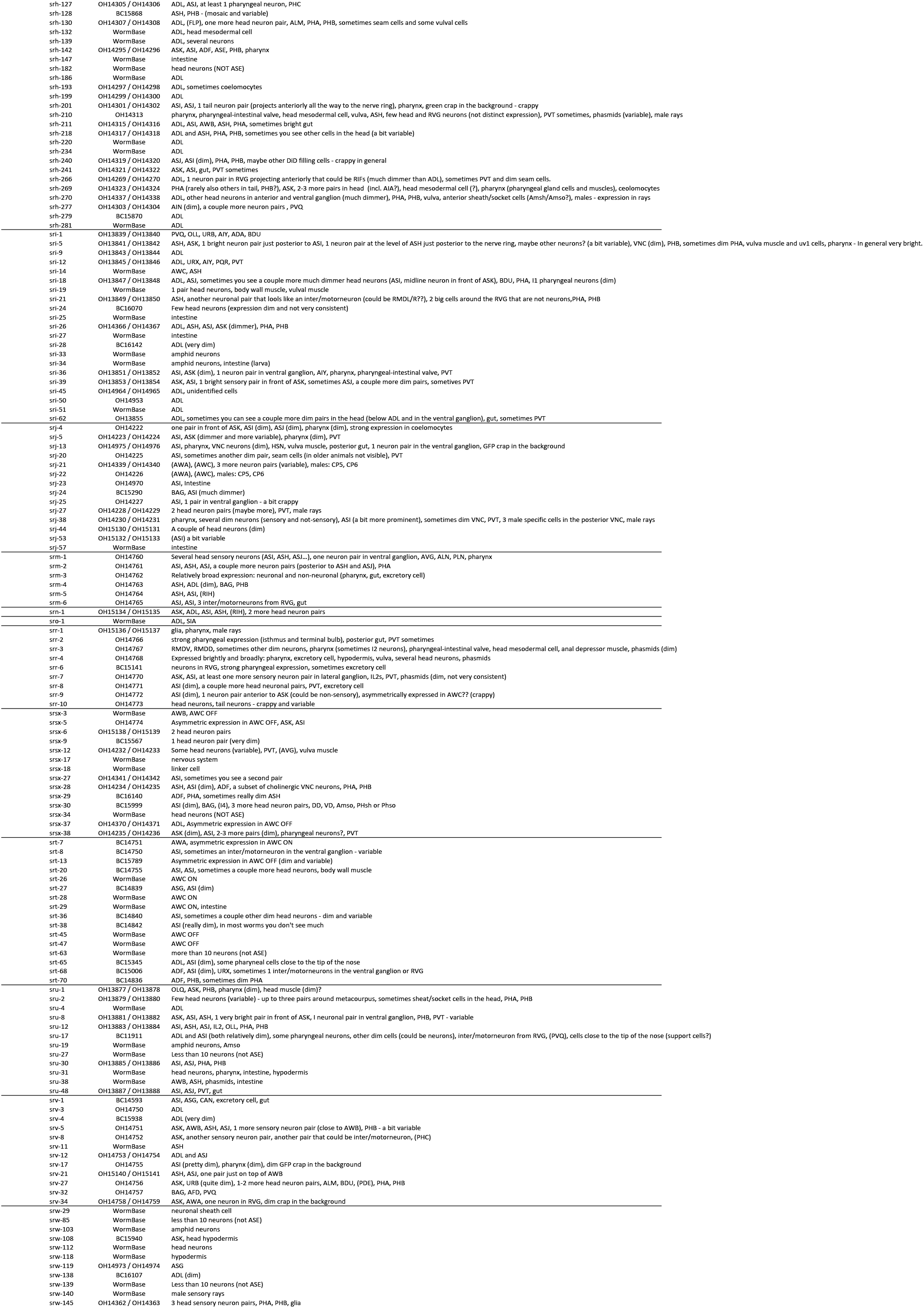
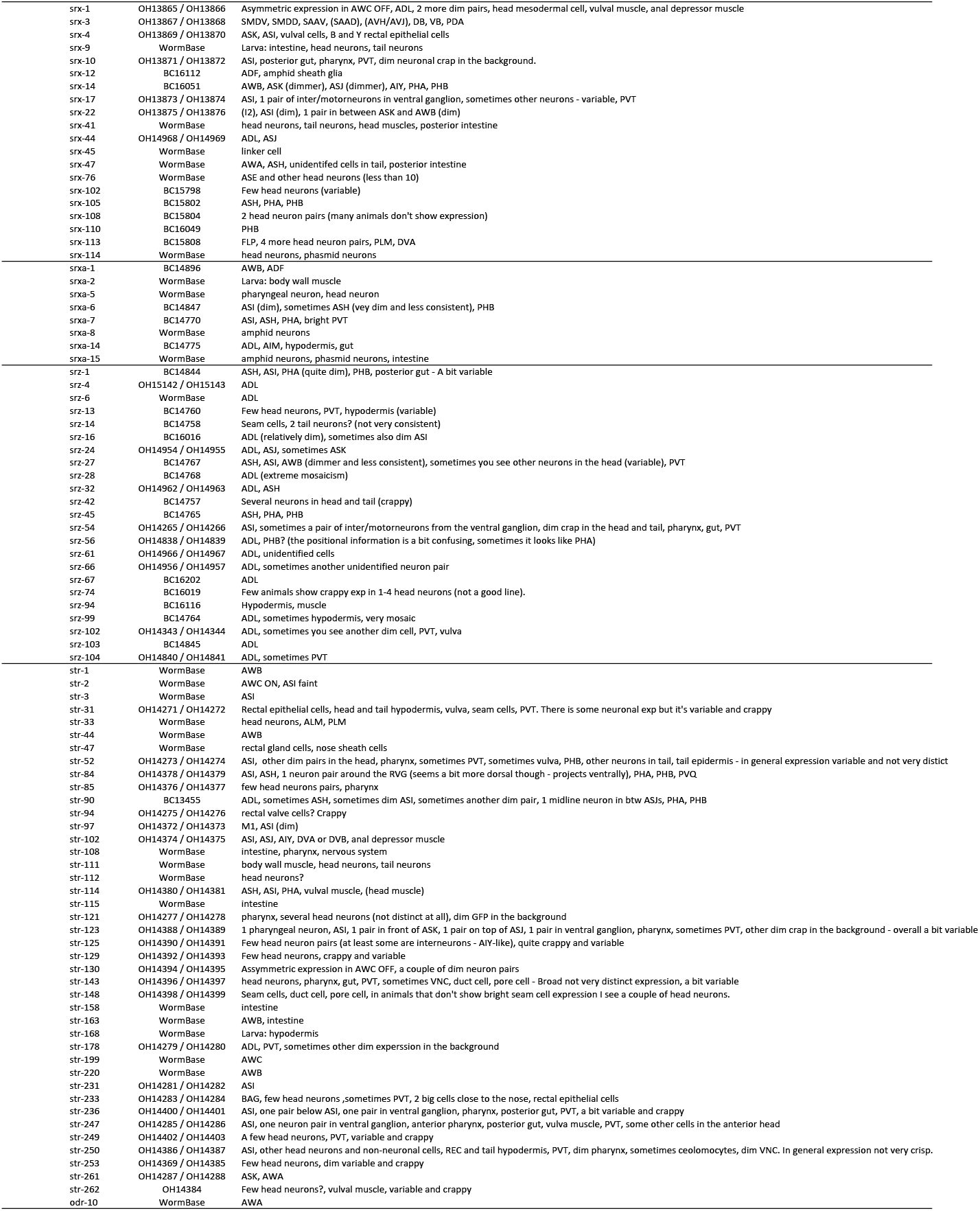
Masterlist of all examined GPCR reporters

**Table S2:**
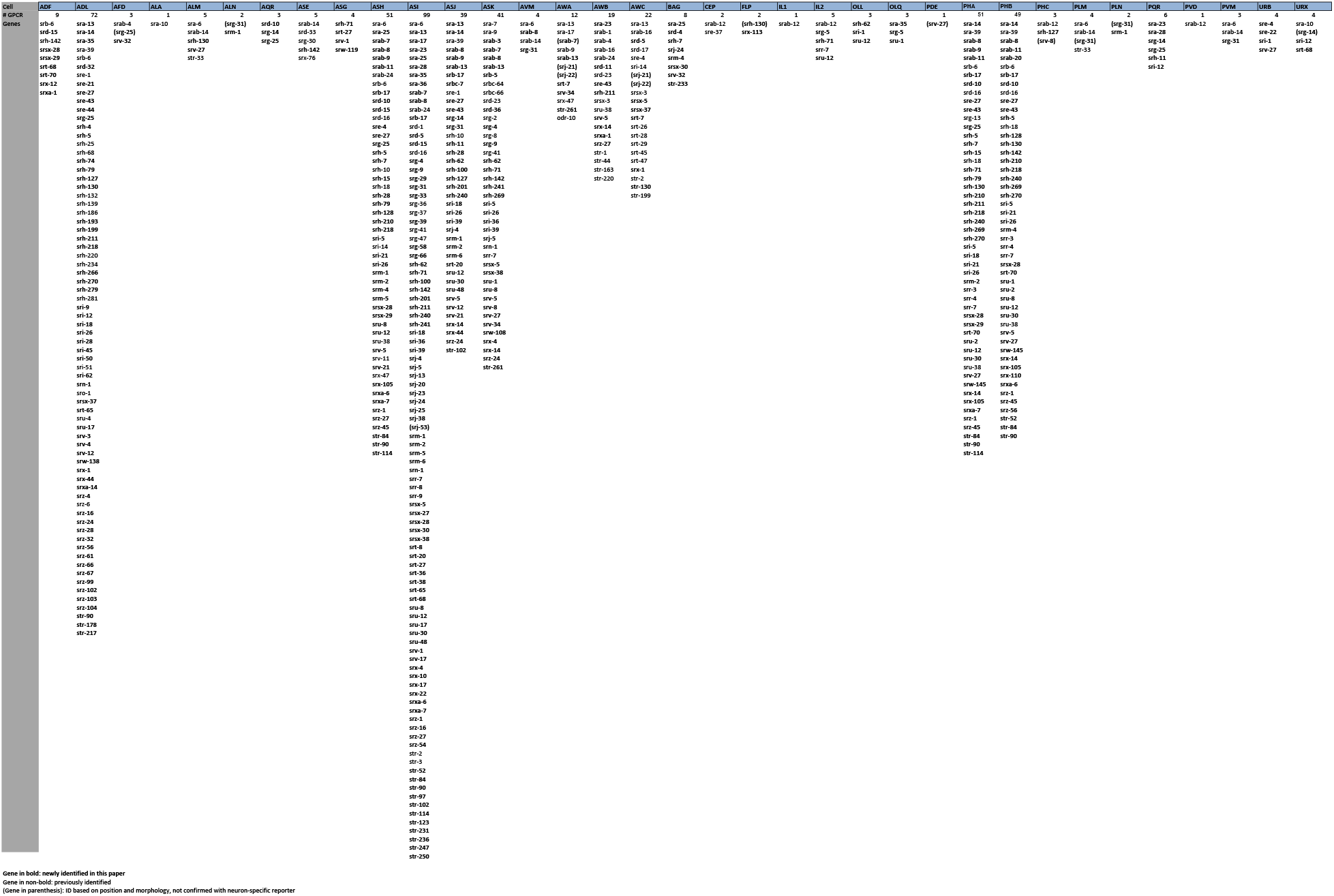
List of all identified sensory neurons with GPCR expression

**Table S3:**
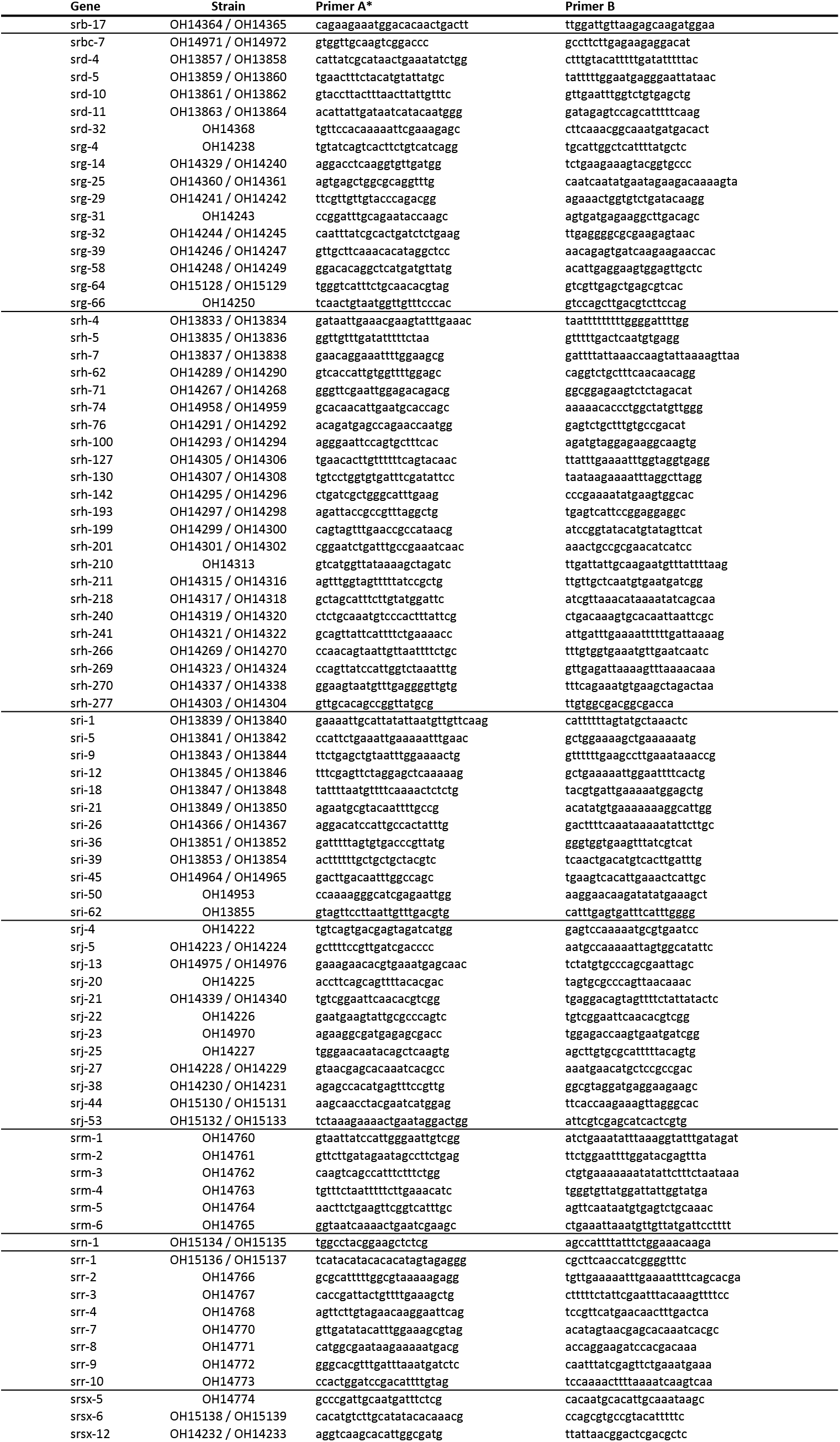
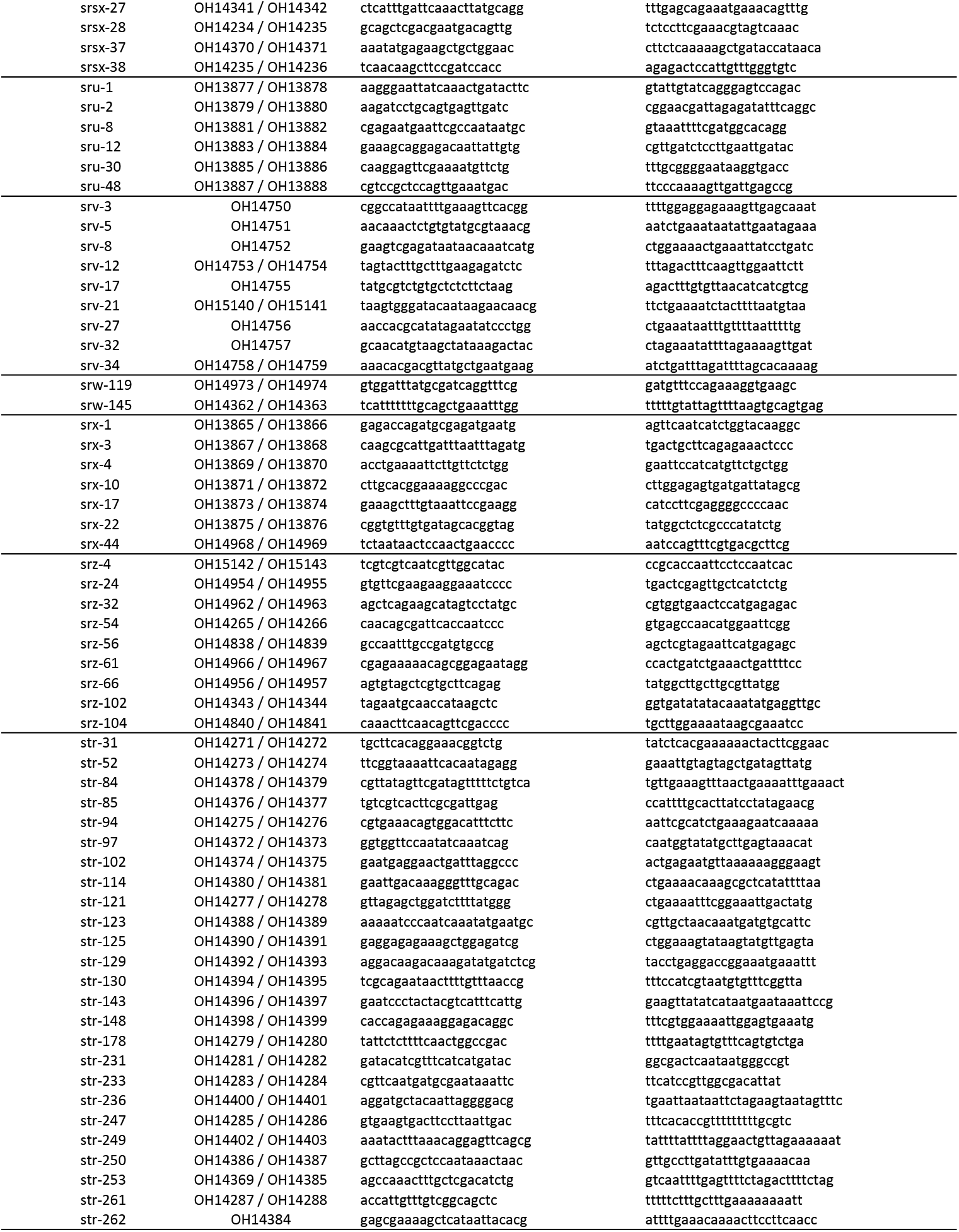
Primers. Primer sequences for the reporters generated by the Vancouver consortium (BC strains) can be found at http://www.gfpworm.org

## REFERENCES

1. Hobert O, Carrera I, Stefanakis N. The molecular and gene regulatory signature of a neuron. Trends in neurosciences. 2010;33(10):435–45. Epub 2010/07/29. doi: S0166-2236(10)00082-2 [pii] 10.1016/j.tins.2010.05.006. PubMed PMID: 20663572.

2. Troemel ER, Chou JH, Dwyer ND, Colbert HA, Bargmann CI. Divergent seven transmembrane receptors are candidate chemosensory receptors in C. elegans. Cell. 1995;83(2):207–18.

3. Chalfie M, Tu Y, Euskirchen G, Ward WW, Prasher DC. Green fluorescent protein as a marker for gene expression. Science. 1994;263(5148):802–5. PubMed PMID: 8303295.

4. Lagerstrom MC, Schioth HB. Structural diversity of G protein-coupled receptors and significance for drug discovery. Nat Rev Drug Discov. 2008;7(4):339–57. Epub 2008/04/03. doi: nrd2518 [pii] 10.1038/nrd2518. PubMed PMID: 18382464.

5. Fredriksson R, Lagerstrom MC, Lundin LG, Schioth HB. The G-protein-coupled receptors in the human genome form five main families. Phylogenetic analysis, paralogon groups, and fingerprints. Mol Pharmacol. 2003;63(6):1256–72. doi: 10.1124/mol.63.6.1256. PubMed PMID: 12761335.

6. Foster SR, Roura E, Thomas WG. Extrasensory perception: odorant and taste receptors beyond the nose and mouth. Pharmacol Ther. 2014;142(1):41–61. doi: 10.1016/j.pharmthera.2013.11.004. PubMed PMID: 24280065.

7. Thomas JH, Robertson HM. The Caenorhabditis chemoreceptor gene families. BMC Biol. 2008;6:42. doi: 10.1186/1741-7007-6-42. PubMed PMID: 18837995; PubMed Central PMCID: PMC2576165.

8. The-C.elegans-Sequencing-Consortium. Genome sequence of the nematode C. elegans: a platform for investigating biology. Science. 1998;282(5396):2012–8.

9. White JG, Southgate E, Thomson JN, Brenner S. The structure of the nervous system of the nematode *Caenorhabditis elegans*. Philosophical Transactions of the Royal Society of London B Biological Sciences. 1986;314:1–340.

10. Hobert O. The neuronal genome of Caenorhabditis elegans. WormBook. 2013:1–106. doi: 10.1895/wormbook.1.161.1. PubMed PMID: 24081909.

11. Robertson HM, Thomas JH. The putative chemoreceptor families of C. elegans. WormBook. 2006:1–12. Epub 2007/12/01. doi: 10.1895/wormbook.1.66.1. PubMed PMID: 18050473.

12. Taniguchi G, Uozumi T, Kiriyama K, Kamizaki T, Hirotsu T. Screening of odor-receptor pairs in Caenorhabditis elegans reveals different receptors for high and low odor concentrations. Sci Signal. 2014;7(323):ra39. doi: 10.1126/scisignal.2005136. PubMed PMID: 24782565.

13. Sengupta P, Chou JH, Bargmann CI. odr-10 encodes a seven transmembrane domain olfactory receptor required for responses to the odorant diacetyl. Cell. 1996;84(6):899–909. Epub 1996/03/22. doi: S0092-8674(00)81068-5 [pii]. PubMed PMID: 8601313.

14. Zhang Y, Chou JH, Bradley J, Bargmann CI, Zinn K. The Caenorhabditis elegans seven-transmembrane protein ODR-10 functions as an odorant receptor in mammalian cells. Proc Natl Acad Sci U S A. 1997;94(22):12162–7. PubMed PMID: 9342380; PubMed Central PMCID: PMCPMC23737.

15. Park D, O’Doherty I, Somvanshi RK, Bethke A, Schroeder FC, Kumar U, et al. Interaction of structure-specific and promiscuous G-protein-coupled receptors mediates small-molecule signaling in Caenorhabditis elegans. Proc Natl Acad Sci U S A. 2012. Epub 2012/06/06. doi: 1202216109 [pii] 10.1073/pnas.1202216109. PubMed PMID: 22665789.

16. McGrath PT, Xu Y, Ailion M, Garrison JL, Butcher RA, Bargmann CI. Parallel evolution of domesticated Caenorhabditis species targets pheromone receptor genes. Nature. 2011; 477(7364):321–5. Epub 2011/08/19. doi: nature10378 [pii] 10.1038/nature10378. PubMed PMID: 21849976; PubMed Central PMCID: PMC3257054.

17. Kim K, Sato K, Shibuya M, Zeiger DM, Butcher RA, Ragains JR, et al. Two chemoreceptors mediate developmental effects of dauer pheromone in C. elegans. Science. 2009;326(5955):994–8. Epub 2009/10/03. doi: 1176331 [pii] 10.1126/science.1176331. PubMed PMID: 19797623.

18. Greene JS, Dobosiewicz M, Butcher RA, McGrath PT, Bargmann CI. Regulatory changes in two chemoreceptor genes contribute to a Caenorhabditis elegans QTL for foraging behavior. eLife. 2016;5. doi: 10.7554/eLife.21454. PubMed PMID: 27893361; PubMed Central PMCID: PMC5125752.

19. Greene JS, Brown M, Dobosiewicz M, Ishida IG, Macosko EZ, Zhang X, et al. Balancing selection shapes density-dependent foraging behaviour. Nature. 2016;539(7628):254–8. doi: 10.1038/nature19848. PubMed PMID: 27799655; PubMed Central PMCID: PMC5161598.

20. Zhang C, Zhao N, Chen Y, Zhang D, Yan J, Zou W, et al. The Signaling Pathway of Caenorhabditis elegans Mediates Chemotaxis Response to the Attractant 2-Heptanone in a Trojan Horse-like Pathogenesis. J Biol Chem. 2016;291(45):23618–27. doi: 10.1074/jbc.M116.741132. PubMed PMID: 27660389; PubMed Central PMCID: PMC5095415.

21. Aoki R, Yagami T, Sasakura H, Ogura K, Kajihara Y, Ibi M, et al. A seven-transmembrane receptor that mediates avoidance response to dihydrocaffeic acid, a water-soluble repellent in Caenorhabditis elegans. J Neurosci. 2011;31(46):16603–10. doi: 10.1523/JNEUROSCI.4018-11.2011. PubMed PMID: 22090488.

22. Peterlin Z, Firestein S, Rogers ME. The state of the art of odorant receptor deorphanization: a report from the orphanage. J Gen Physiol. 2014;143(5):527–42. doi: 10.1085/jgp.201311151. PubMed PMID: 24733839; PubMed Central PMCID: PMCPMC4003190.

23. Montagne N, de Fouchier A, Newcomb RD, Jacquin-Joly E. Advances in the identification and characterization of olfactory receptors in insects. Prog Mol Biol Transl Sci. 2015;130:55–80. doi: 10.1016/bs.pmbts.2014.11.003. PubMed PMID: 25623337.

24. Chen N, Pai S, Zhao Z, Mah A, Newbury R, Johnsen RC, et al. Identification of a nematode chemosensory gene family. Proc Natl Acad Sci U S A. 2005;102(1):146–51. PubMed PMID: 15618405.

25. Wenick AS, Hobert O. Genomic cis-Regulatory Architecture and trans-Acting Regulators of a Single Interneuron-Specific Gene Battery in C. elegans. Dev Cell. 2004;6(6):757–70. PubMed PMID: 15177025.

26. Battu G, Hoier EF, Hajnal A. The C. elegans G-protein-coupled receptor SRA-13 inhibits RAS/MAPK signalling during olfaction and vulval development. Development. 2003; 130(12):2567–77. PubMed PMID: 12736202.

27. Colosimo ME, Brown A, Mukhopadhyay S, Gabel C, Lanjuin AE, Samuel AD, et al. Identification of thermosensory and olfactory neuron-specific genes via expression profiling of single neuron types. Curr Biol. 2004;14(24):2245–51. PubMed PMID: 15620651.

28. McCarroll SA, Li H, Bargmann CI. Identification of transcriptional regulatory elements in chemosensory receptor genes by probabilistic segmentation. Curr Biol. 2005;15(4):347–52. doi: 10.1016/j.cub.2005.02.023. PubMed PMID: 15723796.

29. Krishnan A, Almen MS, Fredriksson R, Schioth HB. Insights into the origin of nematode chemosensory GPCRs: putative orthologs of the Srw family are found across several phyla of protostomes. PLoS One. 2014;9(3):e93048. doi: 10.1371/journal.pone.0093048. PubMed PMID: 24663674; PubMed Central PMCID: PMCPMC3963977.

30. Remy JJ, Hobert O. An interneuronal chemoreceptor required for olfactory imprinting in C. elegans. Science. 2005;309(5735):787–90. PubMed PMID: 16051801.

31. Troemel ER, Sagasti A, Bargmann CI. Lateral signaling mediated by axon contact and calcium entry regulates asymmetric odorant receptor expression in C. elegans. Cell. 1999;99(4):387–98.

32. Wes PD, Bargmann CI. C. elegans odour discrimination requires asymmetric diversity in olfactory neurons. Nature. 2001;410(6829):698–701. PubMed PMID: 11287957.

33. Hsieh YW, Alqadah A, Chuang CF. Asymmetric neural development in the Caenorhabditis elegans olfactory system. Genesis. 2014;52(6):544–54. doi: 10.1002/dvg.22744. PubMed PMID: 24478264; PubMed Central Central PMCID: PMC4065219.

34. Peckol EL, Troemel ER, Bargmann CI. Sensory experience and sensory activity regulate chemosensory receptor gene expression in Caenorhabditis elegans. Proc Natl Acad Sci U S A. 2001;98(20): 11032–8.

35. Gruner M, Nelson D, Winbush A, Hintz R, Ryu L, Chung SH, et al. Feeding state, insulin and NPR-1 modulate chemoreceptor gene expression via integration of sensory and circuit inputs. PLoS Genet. 2014;10(10):e1004707. doi: 10.1371/journal.pgen.1004707. PubMed PMID: 25357003; PubMed Central PMCID: PMCPMC4214617.

36. Nolan KM, Sarafi-Reinach TR, Horne JG, Saffer AM, Sengupta P. The DAF-7 TGF-beta signaling pathway regulates chemosensory receptor gene expression in C. elegans. Genes Dev. 2002;16(23):3061–73. PubMed PMID: 12464635.

37. Ryan DA, Miller RM, Lee K, Neal SJ, Fagan KA, Sengupta P, et al. Sex, age, and hunger regulate behavioral prioritization through dynamic modulation of chemoreceptor expression. Curr Biol. 2014;24(21):2509–17. doi: 10.1016/j.cub.2014.09.032. PubMed PMID: 25438941; PubMed Central PMCID: PMCPMC4214617.

38. Portman DS. Genetic control of sex differences in C. elegans neurobiology and behavior. Adv Genet. 2007;59:1–37. doi: 10.1016/S0065-2660(07)59001-2. PubMed PMID: 17888793.

39. Brenner S. The genetics of Caenorhabditis elegans. Genetics. 1974;77(1):71–94.

40. Granato M, Schnabel H, Schnabel R. pha-1, a selectable marker for gene transfer in C. elegans. Nucleic Acids Res. 1994;22(9):1762–3. PubMed PMID: 8202383; PubMed Central PMCID: PMCPMC4214617.

41. Hodgkin J, Horvitz HR, Brenner S. Nondisjunction Mutants of the Nematode CAENORHABDITIS ELEGANS. Genetics. 1979;91(1):67–94. PubMed PMID: 17248881; PubMed Central PMCID: PMCPMC4214617.

42. Reiner DJ, Newton EM, Tian H, Thomas JH. Diverse behavioural defects caused by mutations in Caenorhabditis elegans unc-43 CaM kinase II. Nature. 1999;402(6758): 199–203. doi: 10.1038/46072. PubMed PMID: 10647014.

43. Park EC, Horvitz HR. C. elegans unc-105 mutations affect muscle and are suppressed by other mutations that affect muscle. Genetics. 1986;113(4):853–67. PubMed PMID: 3744029; PubMed Central PMCID: PMCPMC4214617.

44. Chuang CF, Vanhoven MK, Fetter RD, Verselis VK, Bargmann CI. An innexin-dependent cell network establishes left-right neuronal asymmetry in C. elegans. Cell. 2007; 129(4):787–99. Epub 2007/05/22. doi: S0092-8674(07)00451-5 [pii] 10.1016/j.cell.2007.02.052. PubMed PMID: 17512411.

45. Hobert O. PCR fusion-based approach to create reporter gene constructs for expression analysis in transgenic C. elegans. BioTechniques. 2002;32(4):728–30. PubMed PMID: 11962590.

46. Hunt-Newbury R, Viveiros R, Johnsen R, Mah A, Anastas D, Fang L, et al. High-throughput in vivo analysis of gene expression in Caenorhabditis elegans. PLoS Biol. 2007;5(9):e237. Epub 2007/09/14. doi: 07-PLBI-RA-0103 [pii] 10.1371/journal.pbio.0050237. PubMed PMID: 17850180; PubMed Central Central PMCID: PMC4065219.

47. Schindelin J, Arganda-Carreras I, Frise E, Kaynig V, Longair M, Pietzsch T, et al. Fiji: an open-source platform for biological-image analysis. Nat Methods. 2012;9(7):676–82. doi: 10.1038/nmeth.2019. PubMed PMID: 22743772; PubMed Central PMCID: PMCPMC4214617.

48. Chang C, Hsieh YW, Lesch BJ, Bargmann CI, Chuang CF. Microtubule-based localization of a synaptic calcium-signaling complex is required for left-right neuronal asymmetry in C. elegans. Development. 2011;138(16):3509–18. doi: 10.1242/dev.069740. PubMed PMID: 21771813; PubMed Central Central PMCID: PMC4065219.

49. Serrano-Saiz E, Oren-Suissa M, Bayer EA, Hobert O. Sexually Dimorphic Differentiation of a C. elegans Hub Neuron Is Cell Autonomously Controlled by a Conserved Transcription Factor. Curr Biol. 2017;27(2):199–209. doi: 10.1016/j.cub.2016.11.045. PubMed PMID: 28065609.

50. Pereira L, Kratsios P, Serrano-Saiz E, Sheftel H, Mayo AE, Hall DH, et al. A cellular and regulatory map of the cholinergic nervous system of C. elegans. eLife. 2015;4. doi: 10.7554/eLife.12432. PubMed PMID: 26705699.

51. Gendrel M, Atlas EG, Hobert O. A cellular and regulatory map of the GABAergic nervous system of C. elegans. eLife. 2016;5. doi: 10.7554/eLife.17686. PubMed PMID: 27740909; PubMed Central Central PMCID: PMC4065219.

52. Howe KL, Bolt BJ, Cain S, Chan J, Chen WJ, Davis P, et al. WormBase 2016: expanding to enable helminth genomic research. Nucleic Acids Res. 2016;44(D1):D774–80. doi: 10.1093/nar/gkv1217. PubMed PMID: 26578572; PubMed Central Central PMCID: PMC4065219.

53. Shimodaira H. Approximately unbiased tests of regions using multistep-multiscale bootstrap resampling. The Annals of Statistics. 2004;32(6):2616–41.

54. Boulin T, Etchberger JF, Hobert O. Reporter gene fusions. WormBook. 2006:1–23. Epub 2007/12/01. doi: 10.1895/wormbook.1.106.1. PubMed PMID: 18050449.

55. Hilliard MA, Bargmann CI, Bazzicalupo P. C. elegans responds to chemical repellents by integrating sensory inputs from the head and the tail. Curr Biol. 2002;12(9):730–4. PubMed PMID: 12007416.

56. Zou W, Cheng H, Li S, Yue X, Xue Y, Chen S, et al. Polymodal Responses in C. elegans Phasmid Neurons Rely on Multiple Intracellular and Intercellular Signaling Pathways. Scientific reports. 2017;7:42295. doi: 10.1038/srep42295. PubMed PMID: 28195191; PubMed Central PMCID: PMCPMC4214617.

57. Micallef L, Rodgers P. eulerAPE: drawing area-proportional 3-Venn diagrams using ellipses. PLoS One. 2014;9(7):e101717. doi: 10.1371/journal.pone.0101717. PubMed PMID: 25032825; PubMed Central Central PMCID: PMC4065219.

58. Hobert O, Johnston RJ, Jr., Chang S. Left-right asymmetry in the nervous system: the Caenorhabditis elegans model. Nat Rev Neurosci. 2002;3(8):629–40. PubMed PMID: 12154364.

59. Yu S, Avery L, Baude E, Garbers DL. Guanylyl cyclase expression in specific sensory neurons: a new family of chemosensory receptors. Proc Natl Acad Sci U S A. 1997;94(7):3384–7.

60. Ortiz CO, Faumont S, Takayama J, Ahmed HK, Goldsmith AD, Pocock R, et al. Lateralized gustatory behavior of C. elegans is controlled by specific receptor-type guanylyl cyclases. Curr Biol. 2009;19(12):996–1004. Epub 2009/06/16. doi: S0960-9822(09)01178-6 [pii] 10.1016/j.cub.2009.05.043. PubMed PMID: 19523832; PubMed Central Central PMCID: PMC4065219.

61. Pierce-Shimomura JT, Morse TM, Lockery SR. The fundamental role of pirouettes in Caenorhabditis elegans chemotaxis. J Neurosci. 1999;19(21):9557–69. PubMed PMID: 10531458.

62. Lesch BJ, Gehrke AR, Bulyk ML, Bargmann CI. Transcriptional regulation and stabilization of left-right neuronal identity in C. elegans. Genes Dev. 2009;23(3):345–58. Epub 2009/02/11. doi: 23/3/345 [pii] 10.1101/gad.1763509. PubMed PMID: 19204119.

63. Cochella L, Tursun B, Hsieh YW, Galindo S, Johnston RJ, Chuang CF, et al. Two distinct types of neuronal asymmetries are controlled by the Caenorhabditis elegans zinc finger transcription factor die-1. Genes Dev. 2014;28(1):34–43. doi: 10.1101/gad.233643.113. PubMed PMID: 24361693.

64. Emmons SW. The development of sexual dimorphism: studies of the Caenorhabditis elegans male. Wiley interdisciplinary reviews Developmental biology. 2014;3(4):239–62. doi: 10.1002/wdev.136. PubMed PMID: 25262817; PubMed Central Central PMCID: PMC4065219.

65. Jarrell TA, Wang Y, Bloniarz AE, Brittin CA, Xu M, Thomson JN, et al. The connectome of a decision-making neural network. Science. 2012;337(6093):437–44. Epub 2012/07/28. doi: 10.1126/science.1221762337/6093/437 [pii]. PubMed PMID: 22837521.

66. Lipton J, Kleemann G, Ghosh R, Lints R, Emmons SW. Mate searching in Caenorhabditis elegans: a genetic model for sex drive in a simple invertebrate. J Neurosci. 2004;24(34):7427–34. doi: 10.1523/JNEUR0SCI.1746-04.2004. PubMed PMID: 15329389.

67. Oren-Suissa M, Bayer EA, Hobert O. Sex-specific pruning of neuronal synapses in Caenorhabditis elegans. Nature. 2016;533(7602):206–11. doi: 10.1038/nature17977. PubMed PMID: 27144354; PubMed Central Central PMCID: PMC4065219.

68. Stefanakis N, Carrera I, Hobert O. Regulatory Logic of Pan-Neuronal Gene Expression in C. elegans. Neuron. 2015;87(4):733–50. doi: 10.1016/j.neuron.2015.07.031. PubMed PMID: 26291158; PubMed Central Central PMCID: PMC4065219.

69. Cassada RC, Russell RL. The dauerlarva, a post-embryonic developmental variant of the nematode Caenorhabditis elegans. Dev Biol. 1975;46(2):326–42. Epub 1975/10/01. doi: 0012-1606(75)90109-8 [pii]. PubMed PMID: 1183723.

70. Androwski RJ, Flatt KM, Schroeder NE. Phenotypic plasticity and remodeling in the stress-induced Caenorhabditis elegans dauer. Wiley interdisciplinary reviews Developmental biology. 2017. doi: 10.1002/wdev.278. PubMed PMID: 28544390.

71. Pierce KL, Premont RT, Lefkowitz RJ. Seven-transmembrane receptors. Nat Rev Mol Cell Biol. 2002;3(9):639–50. doi: 10.1038/nrm908. PubMed PMID: 12209124.

72. Hall SE, Beverly M, Russ C, Nusbaum C, Sengupta P. A cellular memory of developmental history generates phenotypic diversity in C. elegans. Curr Biol. 2010;20(2):149–55. Epub 2010/01/19. doi: S0960-9822(09)02050-8 [pii] 10.1016/j.cub.2009.11.035. PubMed PMID: 20079644; PubMed Central Central PMCID: PMC4065219.

73. Sims JR, Ow MC, Nishiguchi MA, Kim K, Sengupta P, Hall SE. Developmental programming modulates olfactory behavior in C. elegans via endogenous RNAi pathways. eLife. 2016;5. doi: 10.7554/eLife.11642. PubMed PMID: 27351255; PubMed Central PMCID: PMCPMC4214617.

74. Hilliard MA, Apicella AJ, Kerr R, Suzuki H, Bazzicalupo P, Schafer WR. In vivo imaging of C. elegans ASH neurons: cellular response and adaptation to chemical repellents. EMBO J. 2005;24(1):63–72. doi: 10.1038/sj.emboj.7600493. PubMed PMID: 15577941; PubMed Central PMCID: PMCPMC4214617.

